# Enhanced feature matching in single-cell proteomics characterizes response to IFN-γ and reveals co-existence of different cell states

**DOI:** 10.1101/2024.01.10.575010

**Authors:** Karl K. Krull, Syed A. Ali, Jeroen Krijgsveld

**Affiliations:** German Cancer Research Center (DKFZ), Heidelberg, Germany; Heidelberg University, Medical Faculty, Heidelberg, Germany

**Author notes:** Correspondence to J.K.

## Abstract

Proteome analysis by data-independent acquisition (DIA) has become a powerful approach to obtain deep proteome coverage, and has gained recent traction for label-free analysis of single cells. However, optimal experimental design for DIA-based single-cell proteomics has not been fully explored, and performance metrics of subsequent data analysis tools remain to be evaluated. Therefore, we here present DIA-ME, a data analysis strategy that exploits the co-analysis of low-input samples with a so-called matching enhancer (ME) of higher input, to increase sensitivity, proteome coverage, and data completeness. We evaluate the matching specificity of DIA-ME by a two-proteome model, and demonstrate that false discovery and false transfer are maintained at low levels when using DIA-NN software, while preserving quantification accuracy. We apply DIA-ME to investigate the proteome response of U-2 OS cells to interferon gamma (IFN-γ) in single cells, and recapitulate the time-resolved induction of IFN-γ response proteins as observed in bulk material. Moreover, we observe co- and anti-correlating patterns of protein expression within the same cell, indicating mutually exclusive protein modules and the co-existence of different cell states. Collectively our data show that DIA-ME is a powerful, scalable, and easy-to-implement strategy for single-cell proteomics.

## Introduction

The increased awareness of biological heterogeneity within cell populations, evidenced by profound differences in gene transcription among individual cells (1–4) has changed our understanding of the confines of what is regarded as the same cell type (5, 6). This phenomenon helped to explain biological transitions as a continuous process rather than a collection of discrete steps and could also have important implications for understanding and treating diseases (7). However, a description only at the transcriptomic level lacks information about post-transcriptional regulation that translate into functional proteomic changes (8–10), therefore requiring techniques for the proteomic investigation of individual cells. While single-cell RNA sequencing has become routinely accessible, single-cell proteomics (SCP) has undergone a more recent but steep development, benefitting from increased sensitivity offered by novel sample preparation workflows and mass spectrometric instrumentation (10–12).

Yet, SCP still suffers from several limitations in proteomic depth and throughput. Since the material from individual cells is scarce, several studies focused on reducing losses during sample preparation by process miniaturization (13, 14) and avoiding surface adsorption (15, 16). As one important example, the SCoPE-MS (single-cell proteomics by mass spectrometry) methodology uses sample multiplexing via TMT-labeling to concomitantly increase analyte concentration and throughput (17, 18). However, recent reports propose that the inclusion of carrier channels in this method can compress reporter ion ratios, jeopardizing the accuracy of protein quantification (19–21). For this reason, label-free approaches have become a popular alternative, where in particular the use of collective precursor isolation in MS via data-independent acquisition (DIA) showed enhanced sampling depth in shorter time compared to data-dependent acquisition (DDA) (22). Nevertheless, DIA produces chimeric MS/MS spectra containing highly convoluted mixtures of simultaneously fragmented precursors and their fragment ions, thus demanding special ways of data analysis. Here, the introduction of diaPASEF was pivotal, enhancing the signal-to-noise ratio by excluding singly charged ions before MS/MS acquisition (23). Still, to cope with the complexity of the resulting spectra, pre-generated libraries comprising unambiguous information on peptides and their fragments remained inevitable (23, 24). Particularly, this poses challenges in SCP as library-based analyses require time-intensive data collection from preceding runs, varying with instruments and projects, and they demand higher sample input even if cell types are scarce. Novel software tools have partly overcome these constraints by direct (i.e. library-free) analysis of DIA spectra, of which Spectronaut (25) and DIA-NN (26) are the most widely used. Although these tools handle data in slightly different manners, e.g. either utilizing a peptide-centric (DIA-NN) or spectrum-centric approach (Spectronaut), both consist of a two-step process, in which the initial assignment is followed by storing the information in an internal library that is subsequently used to re-analyze the data (26–29). This procedure shares a conceptual resemblance with the match-between-runs (MBR) algorithm that is commonly used in DDA approaches, as it aims to recover unidentified features to mitigate missing values across experiments. Since high data completeness is crucial when analyzing large sample cohorts, as is usually the case in single-cell studies, MBR has frequently been applied in DIA workflows (14, 30–36). While this conveniently leads to better data coverage, the quality of matching in DIA data has not been formally addressed, and in particular it has yet to be determined if diminished signal intensity in low-input data leads to false feature matching. In addition, it is unclear how quality of matching depends on the peptide amount used in the donor data set.

To address these issues and establish an optimized workflow for label-free proteomics of single cells, we present DIA-ME, a data analysis strategy that improves proteome coverage and data completeness in low-input DIA data, while preserving FDR-control and enhancing the reliability of feature matching. In this approach, low-input samples of interest are co-analyzed with a set of matching enhancers (MEs), containing a higher peptide amount of the same origin that is used to identify equivalent features in the matching step during DIA data analysis. While spectral libraries are frequently used in SCP to achieve higher sensitivity (14, 34–38), we show that DIA-ME effectively circumvents this requirement by considerably increasing proteomic depth in library-free searches. Here, we establish DIA-ME employing an in-house generated benchmark dataset of a two-species model system to determine optimal size of ME samples and to assure minimal transfer of false identifications. Using injection amounts equivalent to single cells (200 pg), we demonstrate that DIA-ME improves proteome coverage with high quantitative accuracy compared to conventional data analysis, resulting in the characterization of the IFN-γ-responsive proteome that was highly similar to that obtained by bulk analysis. Finally, DIA-ME-assisted analysis of 143 cells revealed oppositely regulated proteome programs, pointing to mutually exclusive protein expression and the co-existence of different cell states within the same population.

## Results

### The principle of enhanced matching in DIA-ME

A key principle to achieve high data completeness in DIA data is the transfer of peptide information across multiple samples in the same set. This functionality is at the core of Spectronaut (28) and DIA-NN (29), and we endeavored to further exploit its capabilities in low-input proteomic data, which usually suffers from missing values and limited sensitivity. Specifically, both tools handle DIA data in a 2-step procedure (despite fundamental differences in their architecture), storing information on precursor and fragment ions in an internal spectral library after the initial identification step, which is then used in a second pass to specifically extract elution profiles from other runs that are analyzed in parallel. This last step considerably reduces missing values across samples, and in the following we refer to it as “MBR” (match-between-runs), although this has been originally coined for DDA data. We speculated that expanding the resources of the internal library could help to concomitantly improve sensitivity and data completeness of proteomic experiments, especially when data are sparse as in single-cell analyses. To this end, we conceived the DIA-ME concept, wherein files with low-input runs are jointly analyzed with runs from higher sample input, effectively serving as a reference database. We termed the files from higher input samples “matching enhancers” (MEs), as they contribute to the creation of an enlarged internal library that is used to extract similar signals from low-input runs, and thus improve sensitivity of the analysis (**Fig. 1A**). DIA-ME is easy to implement in existing proteomic workflows, as it only requires running a few MEs together with a series of low-input samples, followed by analyzing the collective data in existing software tools (**Fig. 1B**). In this work, we aimed to critically assess the performance of DIA-ME, and apply it to the investigation of single cells.

**Figure 1:**
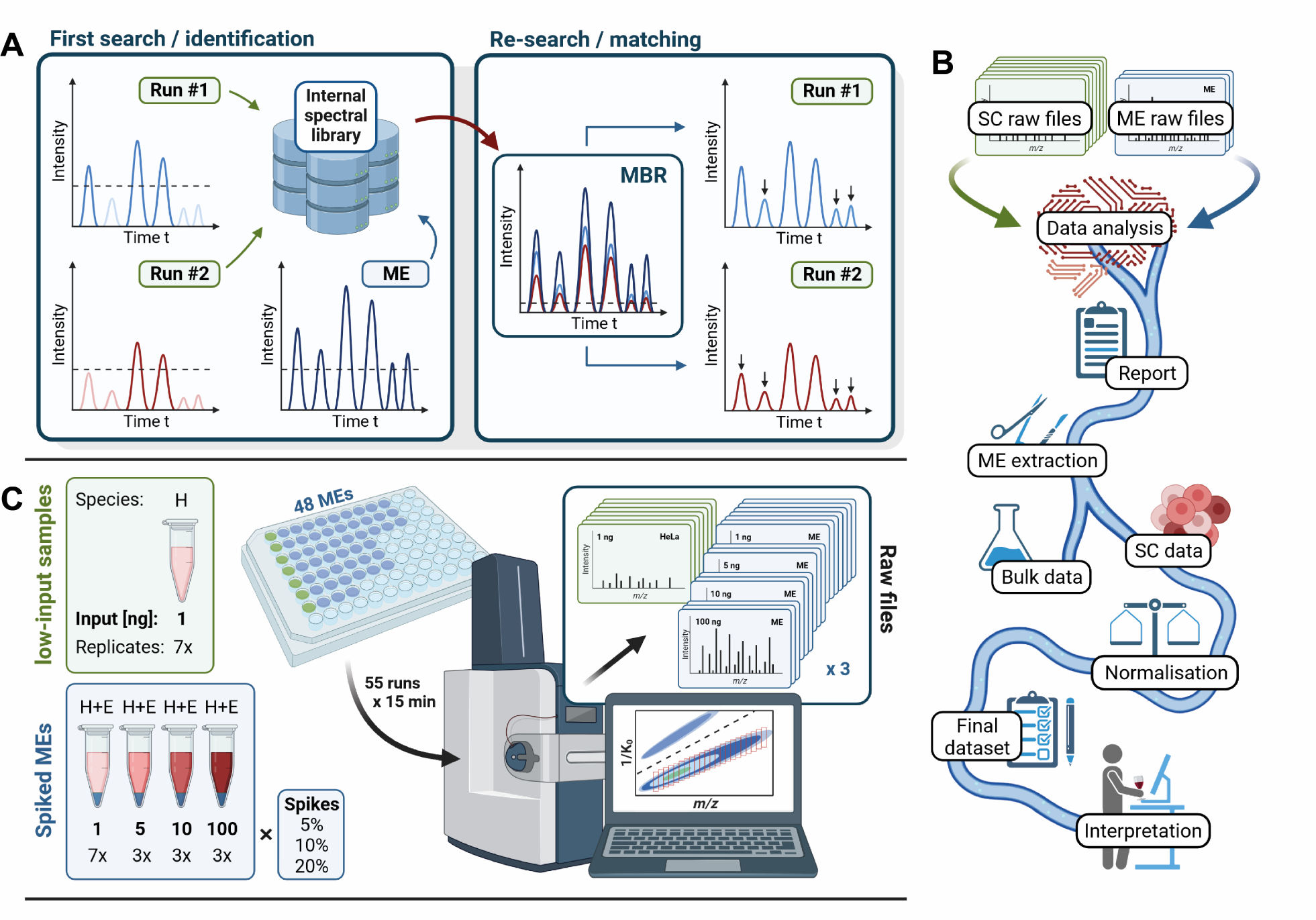
Concept of DIA-ME – an analytical workflow for single-cell proteomics. (A) The principle of DIA-ME: DIA data analysis by DIA-NN and Spectronaut is based on a two-step process, in which peptides are identified and subsequently gathered in an internal library, before the stored information is used to re-search the data. By providing a higher-input sample (Matching enhancer: ME) to the first search, more information can be gathered in the library to help identifying low-abundance signals in the low-input runs. (B) Data pipeline for implementing DIA-ME: low-input samples of interest (e.g. single-cell (SC) data) are submitted to a DIA data analysis software together with a small number of MEs, while inter-sample matching by MBR is permitted (settings not shown). The resulting report file contains all identified peptides, including those from ME samples. The latter information is removed (drop respective columns/rows) to allow suitable peptide intensity normalization. (C) Two-species model experiment to evaluate data analysis by DIA-ME. Two types of samples were prepared: seven low-input replicates containing an H.sapiens proteome (HeLa cells) (H, green) and twelve sets of H.sapiens samples spiked with E.coli K12 proteome (E, blue). The proteomic mixtures differed in their spiking ratio (5 – 20%) and in their total peptide amount (1 – 100 ng). As a result, a total of 55 runs (7 non-spiked and 48 spiked replicates) were acquired on a timsTOF Pro instrument in diaPASEF mode.

### DIA-ME expands proteomic coverage from low-input data

Having conceptualized the DIA-ME workflow, we aimed to assess the gain in proteome coverage that can be achieved, to determine the optimal size of the ME samples, and to evaluate potential false matching events between MEs and low-input samples. To benchmark this, we generated a ME sample set that consists of mixed HeLa and *Escherichia coli* (*E.coli*) peptides, where the former serve as a “donor” for feature matching to a low-input human “acceptor” sample, while the latter introduce features that are used to assess false transfers. Specifically, we prepared samples of HeLa digests spiked with different ratios of *E.coli* peptides (5 – 20%), and adding up to different total peptide amounts of 1 – 100 ng (**Fig. 1C**). For all LC-MS analyses, we used an active LC gradient of 15 min, and performed MS on a Bruker timsTOF Pro instrument in diaPASEF mode (**Fig. S1A**). Data analysis in DIA-NN led to the identification of approx. 8,000 and 3,500 protein groups in 100-ng and 1-ng samples, respectively (**Fig. S1B**). As we intended, *E.coli* peptide intensities increased by more than two orders of magnitude, thereby scaling with the amount of spiked *E.coli* peptides (**Fig. S1C** and **D**). Consequently, our two-proteome samples contained highly variable records of *E.coli* features ranging from around 500 to 6,000 identified peptides, making it a suitable system to evaluate correct matching in low-input data.

Next, we analyzed a set of 1-ng (non-spiked) HeLa peptide samples under the same LC-MS conditions (**Fig. S2**) and investigated the results from different processing methods (**Fig. 2A**, **Table S1** and **S2**). Conventional analysis of this HeLa sample set via DIA-NN identified on average 2,800 proteins, which was increased to 3,300 when applying MBR (2,600 and 3,100 proteins, respectively, when using Spectronaut), confirming the benefit of activating this function (**Fig. 2B**). Remarkably, co-analysis of HeLa with ME samples drastically improved proteome coverage even further, reaching up to approx. 4,650 proteins in 1-ng HeLa samples when analyzed along with 10-ng MEs (i.e. 10x ME) by either DIA-NN or Spectronaut (**Fig. 2B**). This improvement corresponds to an increase of around 60% and 70% in comparison to the initially performed individual analysis in DIA-NN and Spectronaut, respectively, and was primarily caused by the re-extraction of features that were additionally identified from ME files (**Fig. 2C, D** and **Fig. S3**). We ranked protein groups based on their reported intensities and demonstrated that additional ME-derived identifications also considerably contributed to an improved data completeness in DIA-ME analysis (**Fig. 2E**). Dividing the abundance range into equally sized bins, we further revealed that newly identified proteins were found across the entire scale, but mainly helped to reduce missing values in the low intensity area, thereby increasing data completeness in the two lowest bins from 24.9% to 62.4% and from 3.7 to 36.9%, respectively (in comparison to MBR alone). Moreover, DIA-ME-enabled identification of low-abundance proteins led to an expansion of the protein dynamic range by a full order of magnitude (**Fig. 2F**). This observation was accompanied by the consistent quantification of several low-abundance cell cycle-dependent proteins that were not identified using MBR alone. Notably, we recognized that the average number of identified proteins saturated for the DIA-ME analysis at 10-ng (i.e. 10x MEs) in DIA-NN (**Fig. 2B**), which suggests that identifications cannot be increased infinitely, and that identity transfer does not occur randomly (explored in more detail below).

**Figure 2:**
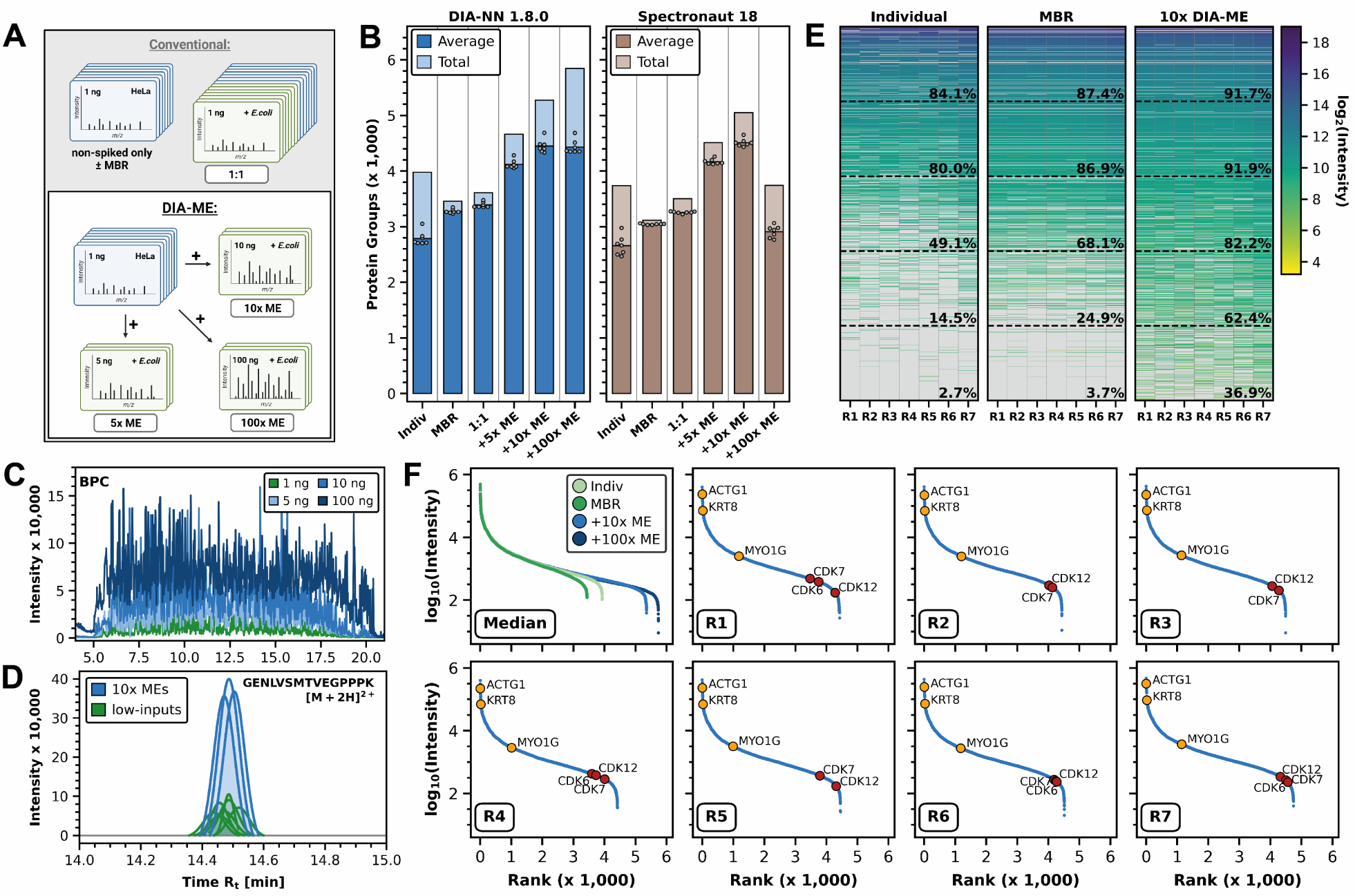
DIA-ME enables ultra-high sensitivity and data completeness. (A) Compositions of raw files for DIA data analysis. In a conventional approach, low-input runs that only contain H.sapiens proteome (blue) were analyzed with or without MBR or in conjunction with *E.coli*-spiked low-input samples of equal injection amount (green). For DIA-ME analysis, low-input *H.sapiens* samples (blue) were analyzed together with a triplicate of 5-ng (5x ME), 10-ng (10x ME) or 100-ng (100x ME) runs that were spiked with *E.coli* proteome (green). Notably, all spiked samples had a spiking ratio of 5%, 10% and 20%. (B) Average (dark) and total (light) protein group identifications in the set of seven non-spiked 1-ng replicates by DIA-NN (blue) and Spectronaut (brown) after different ways of data analysis (see panel A). Results are only shown for analyses that involved a spiking ratio of 10%. (C) Base peak chromatogram (BPC) overlay of one 1-ng, 5-ng, 10-ng and 100-ng replicate, respectively. (D) Superimposed elution peaks of peptide GENLVSMTVEGPPPK2+ from snRNP-B (Small nuclear ribonucleoprotein-associated proteins B and B’) in 1-ng replicates (green) and 10-ng ME samples (blue). (E) Heatmap of ranked protein group (PG) intensities for individual (i.e. without MBR), MBR and 10x DIA-ME analysis of the seven non-spiked 1-ng HeLa replicates. The abundance range was divided into six bins indicated by dashed lines. Percentages reflect the data completeness found in every bin. (F) Rank plot of median protein group quantities in different types of data analysis (upper left), and rank plots of quantified protein groups in non-spiked 1-ng replicates (R1 – 7) after 10x DIA-ME analysis. Consistent quantification of three high- and medium-abundance cytoskeleton proteins (yellow) and three low-abundance cell cycle-related proteins (red) are indicated, the latter being identified exclusively by DIA-ME analysis.

Analysis of the same data by Spectronaut showed very similar trends, although overall identifying fewer proteins (**Fig. 2B**). Here, the most notable observation was the drop in protein identifications when using 100x DIA-ME. We did not further investigate this phenomenon, but assume that feature matching might be impeded in this software due to the large differences in signal intensities. Collectively, our results show that the concept of extensive feature matching in DIA-ME benefits low-input data by decreasing missing values, augmenting sensitivity, and resulting in enhanced proteome coverage.

### DIA-ME improves qualitative reliability in low-input data

Encouraged by these results, we aimed to assess the reliability of protein identification and feature matching in DIA-ME. Since the vast majority of *E.coli* peptides are not shared with the human proteome, we evaluated coverage of species-specific proteomes in non-spiked 1-ng samples before and after they were co-analyzed with *E.coli*-containing ME samples. In this way, we used the *E.coli* peptides in ME samples as a matching resource to estimate erroneous feature assignment (**Table S1** and **S2**).

We first assessed species-specific protein identifications in our previous results and determined the collective false-positive rate (FPR) of *E.coli* proteins (**Fig. 3A**). When analyzing raw files individually, we found reasonably low FPRs of 0.35% and 0.41% in DIA-NN and Spectronaut, respectively. While samples in this analysis did not contain *E.coli* peptides, the number of false positives increased when they were co-analyzed with spiked samples. As a result, FPR reached up to 1.7% in Spectronaut, which corresponds to almost 8.0% FDR (**Fig. 3A** and **Fig. S4A**), in the experiment of equivalent input amounts (i.e. 1:1). Furthermore, Spectronaut’s ability to correctly assign features was dependent on the data provided for matching, since the FPR scaled with the spiking ratio (**Fig. 3A**). In practice, this might lead to problems if the proteome composition differs substantially between experimental conditions, e.g. when comparing different cell types. In contrast, DIA-NN displayed consistently low levels of false-positive identifications. Notably, the application of DIA-ME resulted in maximum FPR of 0.7%, irrespective of the ME ratio (**Fig. 3A**). These observations suggest that the underlying matching process is considerably less controlled in Spectronaut than in DIA-NN, and underscore that joint analysis of low-input and higher peptide amount samples does not compromise protein identification quality in the context of DIA-NN. Moreover, it is supported by the receiver operating characteristics (ROC) curves, which at a specificity cut off at 95% revealed consistently high sensitivity in DIA-NN up to a sample ratio of 10x ME (**Fig. 3B**).

**Figure 3:**
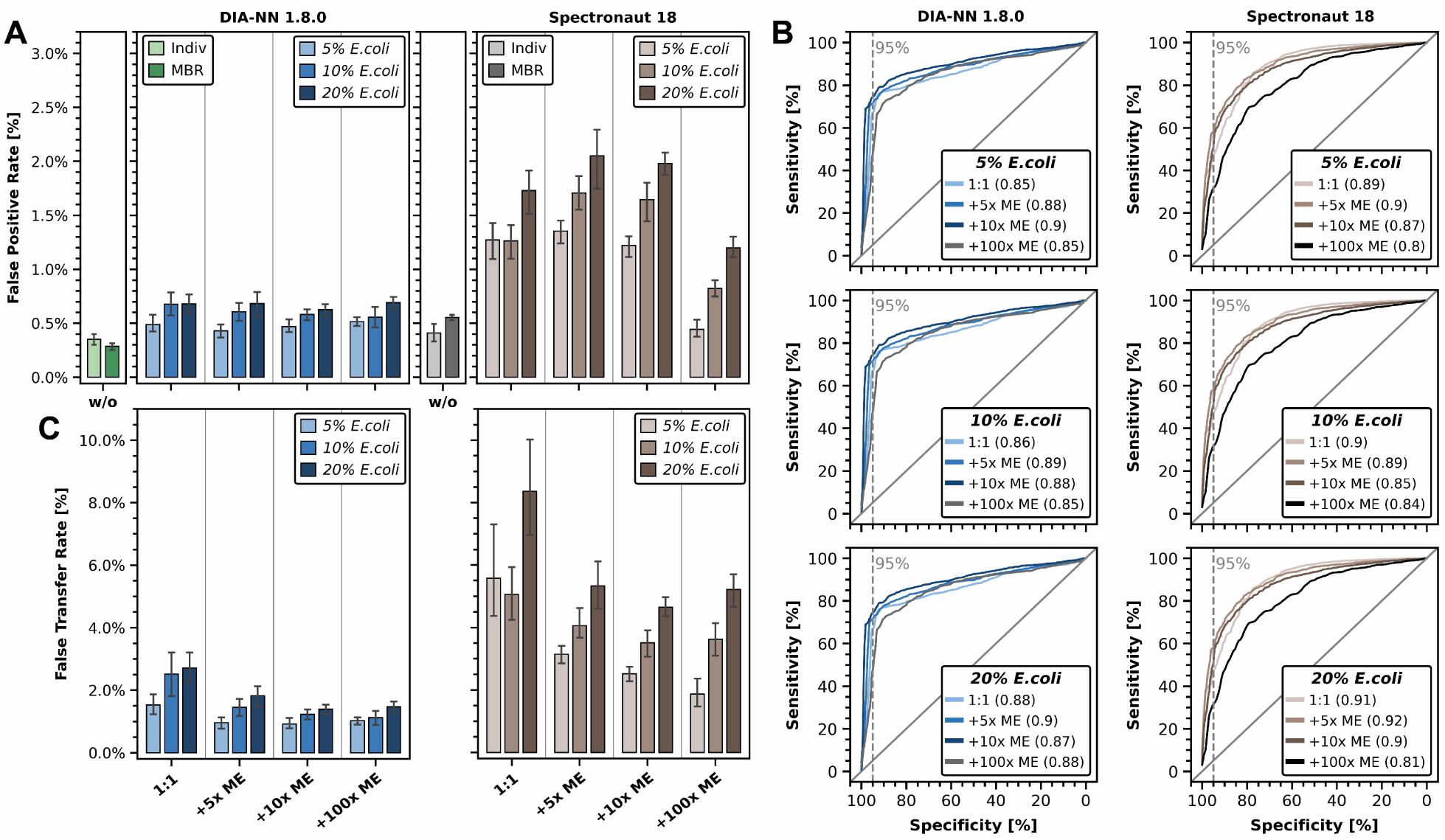
Low FPR and reliable feature matching in DIA-ME. (A) False positive rate, i.e. percentage of detected *E.coli* proteins, in non-spiked 1-ng *H.sapiens* samples for different types of data analysis and DIA software. Analyses without spiked samples are indicated in green and grey (light-: without MBR, dark-: MBR), while analyses together with spiked sample are indicated in blue and brown for DIA-NN and Spectronaut, respectively. The shade of the color indicates the *E.coli* spiking ratio. Error bars are shown as mean ± sd. (B) Receiver operating characteristics (ROC) over data analyses involving spiked ME samples of different injection amounts (DIA-NN: light blue (1:1) to dark grey (100x DIA-ME), Spectronaut: light brown (1:1) to black (100x DIA-ME)) and spiking ratios (top: 5%, middle: 10%, bottom: 20%) for the software DIA-NN (left) and Spectronaut (right) using protein q-value filtering. Points of intersection between the curves and the dashed vertical line represent an observed specificity of 95% in the respective condition. Areas under ROC (AUROC) are indicated in parentheses. (C) False transfer rate, i.e. percentage of *E.coli* proteins among identifications that were transferred by matching, in non-spiked 1-ng *H.sapiens* samples for different types of data analysis and DIA software. Color coding as in panel A. Error bars are shown as mean ± sd.

To estimate the rate of erroneous feature matching in DIA-ME, we next assessed the proportion of *E.coli* proteins among newly identified proteins in HeLa samples after matching (false-transfer rate (FTR)) (**Fig. 3C**). We found elevated FTR when using Spectronaut, incorrectly identifying between 5 – 8% proteins in conventional co-analysis of equal input amounts (i.e. 1:1) (**Fig. 3C**). Contrarily, DIA-NN displayed consistently moderate FTRs of 1.0 – 1.8% across DIA-ME analyses (1.5 – 2.5% for equal input amounts). The observation that this is the case even when matching the 1-ng HeLa sample to 100-ng ME samples that contain 20% *E.coli* peptides indicates high specificity in the matching process that is resilient to the presence of highly abundant interfering peptides. To further examine the specificity of matching, we determined the probability of species-specific proteins that were identified in the MEs and transferred to the low-input samples (transfer rate (TR)) (**Fig. S4B**). In DIA-NN this showed a TR of around 65% for human proteins in comparison to only 5% for *E.coli* proteins in 10x DIA-ME analysis (around 80% and 17%, respectively, in Spectronaut) (**Fig. S4B**), indicating that DIA-NN is superior to Spectronaut in discriminating false- and true-positive identifications during the matching process. Furthermore, in both softwares, we observed a remarkable decline in FTR when employing DIA-ME (**Fig. 3C**). Reduced FTR implies that the better spectral quality in MEs potentially facilitates matching in low-input data when compared to regular MBR among equal inputs. Hence, we not only established that DIA-ME leads to an expanded proteome coverage, but also enhances the fidelity of feature matching in low-input data when co-analyzed with MEs in DIA-NN.

### Precise and accurate quantification with DIA-ME

After successfully showing confident protein identification driven by DIA-ME, we next probed the quality of protein quantification using this concept in DIA-NN. Since normalization is crucial for reliable quantification by compensating for differences between injected peptide amounts, we removed rows from the DIA-NN report that contained peptides identified in ME samples before re-normalizing our datasets. Next, we examined different ways of data normalization for low-input DIA data, including MaxLFQ from the *R* package of DIA-NN (26), *iq* normalization (39) and the recently published directLFQ package in Python (40) (**Fig. S6A**). Remarkably, the output of DIA-NN resulted in insufficient quantitative results for the low-input data, while directLFQ showed the highest accuracy among the tested normalization strategies and it was therefore selected for the following analysis (**Fig. 4**).

**Figure 4:**
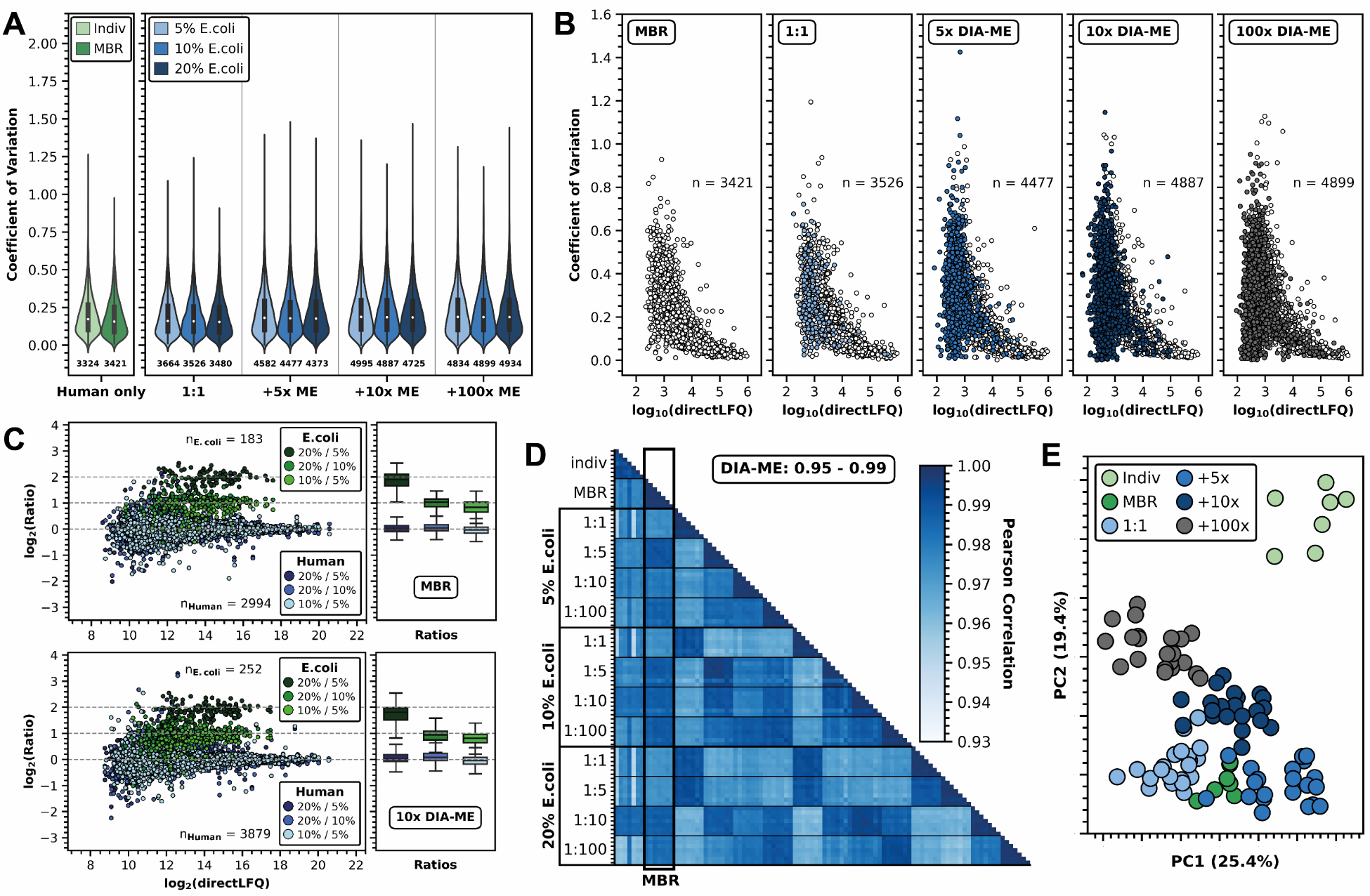
Stable precision and accuracy in protein quantification using directLFQ normalization. (A) Violin plot of human protein CVs in 1-ng samples with (dark green) and without MBR (light green) on the left and after matching against 1-ng (1:1), 5-ng (5x ME), 10-ng (10x ME) and 100-ng (100x ME) samples on the right. Number of proteins in each analysis are indicated beneath the violins, which are colored in blue shades according to the spiking ratio in ME samples. Black boxes in the violins show the dispersion of values between the first and third quartile with the white dot representing the median of the dataset. Box and violin plots showing the entire range from the 0th to 100th percentile. (B) Scatter plot of human protein CV values dependent on their abundance for different types of data analysis (only shown for 10% spiking). White: proteins only identified by MBR; blues and dark grey: proteins exclusively found in DIA-ME analyses. Number of identified proteins per analysis are indicated. (C) Mean protein quantity ratios dependent on their abundance after MBR (top panel) and 10x DIA-ME (bottom panel) analysis, illustrated as scatter plot on the left and box plot on the right. Human and *E.coli* proteins are colored in blues and greens, respectively, with the shade of the color representing the quotient between the different spiking ratios. Expected ratios are depicted as dashed lines. The number of quantified human and *E.coli* proteins are indicated. (D) Pearson correlation heatmap within and among different types of data analysis. Correlations to conventional MBR analysis are framed. The range of observed correlations in DIA- ME analyses is additionally highlighted. (E) Principal component analysis localizing individual replicates from all performed searches in a two-dimensional coordinate system. Data analysis with and without MBR are colored in greens, while DIA-ME analyses using spiked MEs are illustrated in blue shades and dark grey.

For evaluation of the resulting protein quantities, we calculated the coefficient of variation (CV) of protein groups as a measure of the variability within different experiments (**Fig. 4A**). Using directLFQ normalization, we found similar distributions in conventional MBR and DIA-ME analysis with median CV values between 0.16 and 0.19. The slightly higher CVs in DIA-ME experiments may be attributed to the additionally identified proteins, which are generally of lower intensity and are therefore more difficult to quantify precisely (**Fig. 4B**). However, reassuringly, CVs of newly identified proteins scaled very similar with their abundance when compared to proteins that were already identified by conventional MBR, testifying to equivalent consistency in the quantification process. Furthermore, and as anticipated, Pearson correlations of protein quantities across all previously performed experiments showed enhanced quantitative correlation in the MBR approach, surpassing individual analyses (**Fig. 4D**). Interestingly, the application of DIA-ME with varying ME input amounts did not alter this observation and consistently displayed outstanding replicate correlations ranging from 0.95 to 0.99. Moreover, DIA-ME results exhibited strong inter-correlations and aligned with MBR results (indicated by the framed column in **Fig. 4D**). This finding is further substantiated by the results of the principal component analysis (PCA), revealing cohesive clustering of replicates from the same experiment and with results from MBR analysis (**Fig. 4E**). Low quantitative variation and remarkably strong correlation both suggest that highly precise protein quantification is maintained in DIA-ME.

To investigate whether utilization of DIA-ME impacts quantitative accuracy, we assessed if *E.coli* spiking ratios were reflected by relative protein quantification within our spiked 1-ng samples (**Table S3**). Using maxLFQ intensities from DIA-NN, we noticed that protein quantities were estimated inaccurately already when analyzed by conventional MBR (**Fig. S6A**). Moreover, we observed that protein intensities increased uniformly when samples were co-analyzed with higher input ME samples (**Fig. S6B**). Once we used raw peptide intensities and performed protein quantification in the DIA-NN *R* package, this effect was reversed, indicating that it is caused by the internal peptide normalization in DIA-NN, however, did not display accurate quantification (**Fig. S6A**). Re-normalizing peptides by the directLFQ solved both issues and resulted in relative *E.coli* quantities that aligned with their expected ratios >**Fig. 4C, S6A** and **S6C**). This finding emphasizes the importance of normalization for the analysis of low-input DIA data in general, but also for the co-analysis of samples of vastly different input amounts. Hence, we recommend removing peptides that were identified in ME samples from the DIA-NN report before performing directLFQ normalization as a downstream procedure after DIA-ME analysis. Collectively, our data indicate that DIA- ME maintains quantitative accuracy while expanding proteome coverage.

### DIA-ME enables in-depth analysis of single-cell-like input U-2 OS cells upon IFN-γ treatment

Having thoroughly tested the performance of DIA-ME in defined benchmark samples, we next aimed to evaluate its benefits for low-input quantities that are equivalent to single cells, and in particular, to investigate if DIA-ME improves the ability to detect proteome differences between such samples. To this end, we evaluated if DIA-ME can recapitulate the effect in human osteosarcoma cells (U-2 OS) that were treated with Interferon gamma (IFN-γ), an extensively characterized model to study cellular immune response via JAK/STAT signaling (reviewed in (41–43)).

As a starting point, we first generated a reference dataset in a conventional bulk proteomics analysis using 200-ng peptide injections from U-2 OS cells at four time-points during IFN-γ treatment (**Table S4**). Among more than 5,000 identified proteins (**Fig. S7A**) that were quantified with high reproducibility (**Fig. S7B**), we found several proteins to be differentially regulated at distinct time-points (**Fig. S7C**). Yet, the overall effect of IFN-γ on U-2 OS cells was modest, as indicated by the majority of proteins showing minor fold-changes, which did not result in a clear distinction between treated and control samples in a PCA analysis (**Fig. S7D**). Nevertheless, overrepresentation analysis of upregulated proteins showed a clear enrichment of immunologically relevant processes, including the response to IFN-γ (**Fig. S7E**). This was reflected by rapid induction of JAK1 and STAT1 (but not STAT2), and integral parts of the MHC class I pathway, including its building blocks β_2_-microglobulin (B2M) and the corresponding human leukocyte antigens (HLAs) (**Fig. S7H** and **I**). These proteins formed a cluster together with several other known IFN-γ-responding proteins, showing gradual and strong upregulation over 24 hours of treatment (**Fig. S7F** and **G**). For instance, this cluster contained PMSB8, PSMB9 and PSMB10, the three distinguishing members of the IFN- γ-induced immunoproteasome (44–46), as well as TAP1/2 and TAPBP that translocate peptides produced by the immunoproteasome to the endoplasmic reticulum (ER) for loading onto the nascent MHC class I receptor (B2M and HLAs). Consequently, these data show that the proteome response of U-2 OS cells to IFN-γ is mild but detectable in a bulk experiment, identifying known players of the antigen processing and presentation pathway.

We next repeated the IFN-γ treatment experiment, however now using the equivalent of single-cell inputs in a DIA-ME approach. To this end, we collected cells at six time-points during IFN-γ treatment over the course of 24 hours (**Fig. 5A**), using injection amounts of 200 pg per sample, the estimated protein amount of a single U-2 OS cell. For matching purposes, we moreover performed triplicate injections of all six time- points with peptide input amounts of 1, 2 and 10 ng (i.e. 5x, 10x and 50x MEs), which we then co-analyzed with our 200-pg samples using DIA-NN.

**Figure 5:**
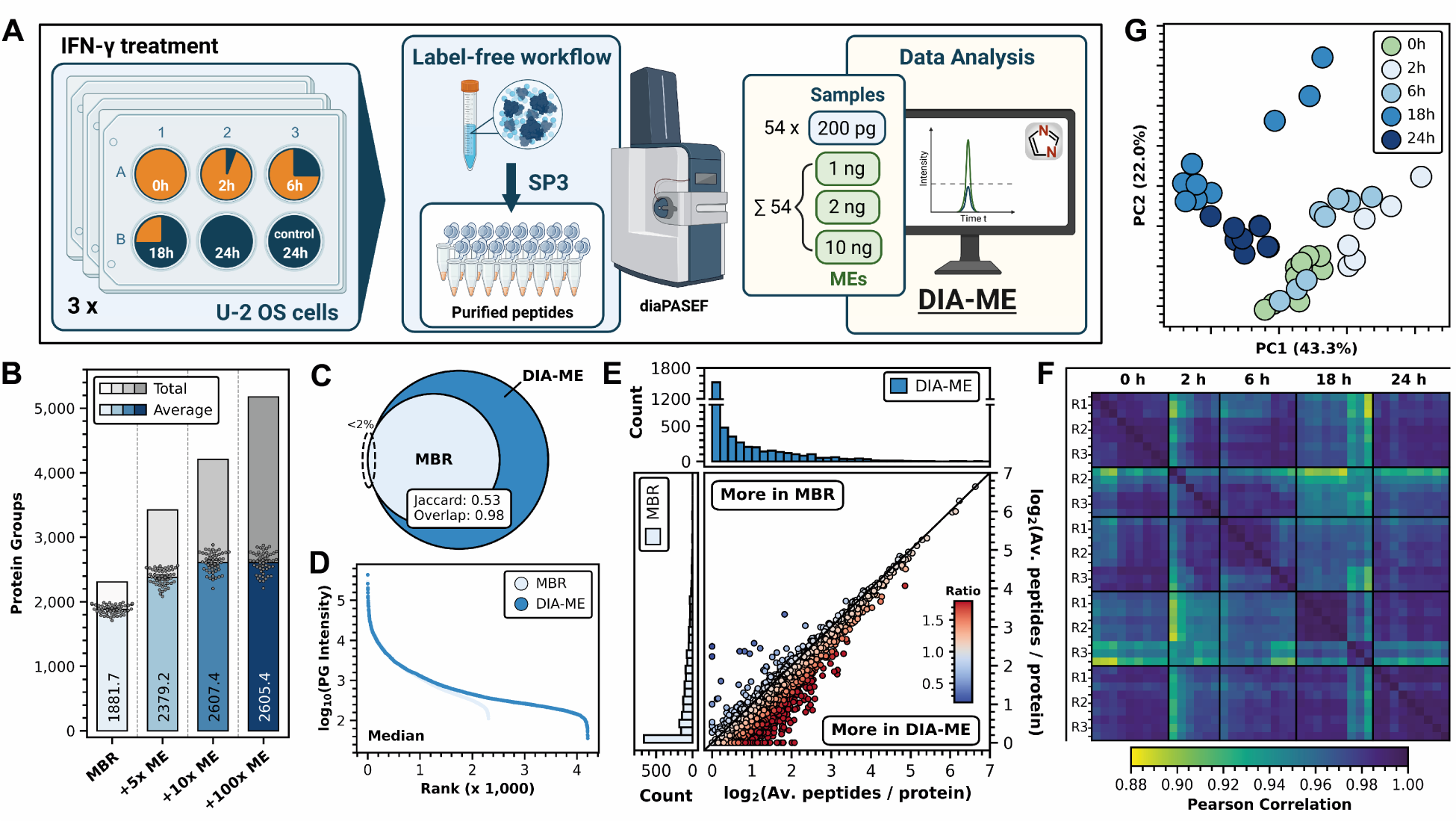
DIA-ME improves proteome coverage in U-2 OS cells upon IFN-γ treatment. (A) Experimental scheme of the IFN-γ treatment and bulk preparation of U-2 OS cells in the DIA-ME workflow. Six time-points from three biological replicates were gathered and peptides diluted to the indicated injection amounts. Three 200-pg injections (blue) and single-injections of 1 – 10 ng (ME samples, green) per time-point and biological replicate made a total of 108 runs. The resulting files were co-analyzed by DIA-ME using DIA-NN. (B) Total (greys) and average identifications of protein groups (blues) in 200-pg samples after DIA-ME analysis with 1 ng (5x ME), 2 ng (10x ME) and 10 ng (50x ME) in comparison to conventional MBR analysis without references samples. Individual number of protein identifications per sample are represented by dark grey dots. MEs were taken from control samples before treatment (0 hours) and from treated samples after 24 hours. (C) Venn diagram of identified protein groups in MBR (light blue) and 10x DIA-ME analysis (blue). The similarity of both populations is described by the overlap (Szymkiewicz– Simpson) and Jaccard index (see method section). (D) Rank plot of median protein group intensities. (E) Joint plot of histogram distributions (top and left side) of average peptide number per protein identified in MBR (y-axis, light blue) and DIA-ME analysis (x-axis, blue). Proteins are illustrated as scatter with their color representing the ratio of peptide identifications between the two analyses, effectively indicating higher coverage in MBR (blue) or in DIA-ME (red). The black line shows peptide number equality (ratio of 1). (F) Pearson correlation heatmap of time-point replicates, showing high correlation in dark blue and lower correlation as green/yellow according to the indicated color bar. (G) Principal component analysis of time-point samples after 10x DIA-ME analysis. Control samples before the treatment are shown in light green while treated samples are illustrated in blue colors.

We used the two extreme time-points (0 h + 24 h) as matching resource for the following analysis (**Table S5**), and increased the average number of identified proteins per 200-pg sample from 1,882 in conventional MBR to 2,607 using DIA-ME (+38%) (**Fig. 5B**), while using other ME time-point samples did not further improve this result (**Fig. S8A**). Interestingly, this was achieved with 10x ME samples and did not further increase with 50x MEs. Since we similarly observed saturation with the 10x ME samples in the previous experiment (**Fig. 2B**), we conclude that the optimal amount of MEs is ratio-dependent rather than being determined by the absolute protein amount in the MEs. Analysis of 200-pg samples with 10x DIA-ME comprised a total of 4,239 protein identifications (**Fig. 5B**), among which 75% were found in at least one replicate per time-point (**Fig. S8B**). Here, around 39% of proteins that were already identified by MBR displayed fewer missing values in DIA-ME (**Fig. S8C**), indicating improved proteome coverage and data completeness. The resulting dataset contained more than 98% of identifications that were also found in MBR, while MBR covered only 53% of proteins identified by DIA-ME (**Fig. 5C**), effectively leading to an expansion of the dynamic range (**Fig. 5D**). Crucially, DIA-ME also identified considerably more peptides per protein (**Fig. 5E, S8D** and **S8E**), providing a stronger basis for protein quantification. Indeed, the DIA- ME dataset displayed great quantitative alignment, exhibiting Pearson correlations between 0.88 and 0.99 across time-point replicates (**Fig. 5F**), while detecting better correlation among earlier and later time- points. This observation is also reflected by the identification of two distinct clusters of earlier and later time-points by PCA analysis (**Fig. 5G**), indicating changes in the proteome composition over the course of the experiment. Taken together, these data show that DIA-ME provides enhanced proteome coverage in single cell-like (200 pg) samples, benefiting from ME samples that are as small as two nanogram, while exhibiting highly reproducible quantification.

To reveal the specific effect of IFN-γ and compare the findings to our reference dataset (**Fig. S7**), we investigated differential protein expression at a depth of more than 3,000 proteins per time-point and detected significant up-regulation of multiple proteins after 24 hours (**Fig. 6A**). As in the bulk (200-ng) samples (**Fig. S7C**), this comprised STAT1, showing the induction of JAK/STAT-mediated signal transduction, and the MHC class I molecules HLA-A/B/C and B2M. Remarkably, DIA-ME enabled the quantification of 35.8% proteins in the total dataset and of 25.2% of known IFN-γ responders (increase of around 56% and 34%, respectively) (**Fig. 6A** and **C**), while their consistent quantification resulted in p-values similar to MBR results (**Fig. S8F**).

**Figure 6:**
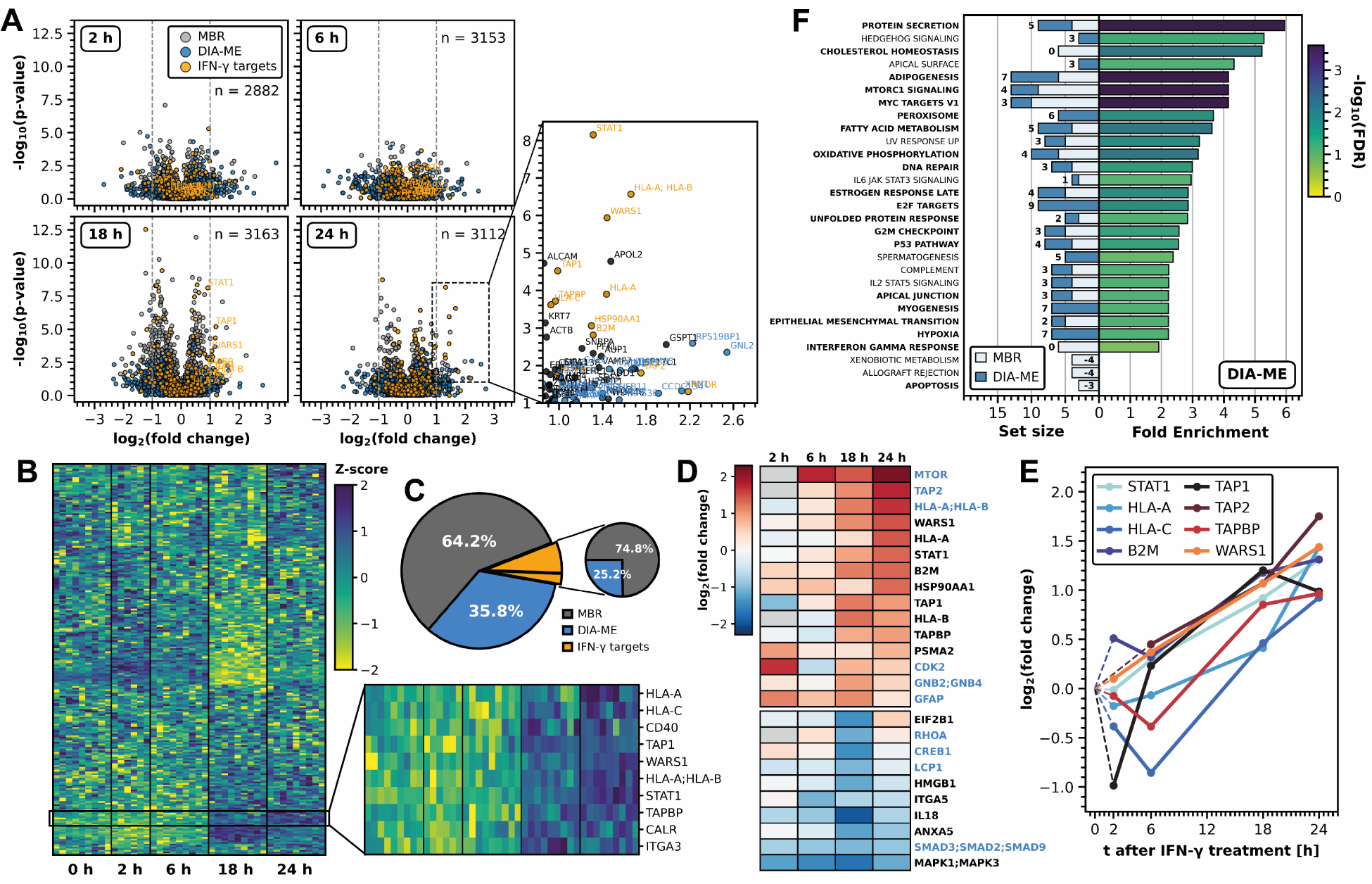
Increased proteomic depth in 200-pg samples allows exploration of IFN-γ-induced immune response in U-2 OS cells. (A) Volcano analysis of Student’s t-test results between 200-pg time-point samples after IFN-γ treatment and the respective 0 hours control without treatment. n: number of differentially expressed proteins. Proteins whose differential expression was observed exclusively after DIA-ME analysis are highlighted in blue, whereas proteins known to be involved in the IFN-γ-response were illustrated in yellow. (B) Heatmap of known IFN-γ-responsive proteins in 200-pg time-point samples after hierarchical clustering by Euclidean distance. Missing values were filled by k-nearest neighbors imputation. Colors illustrate the quantitative changes compared to the protein’s median across all samples by Z-score. The indicated black box outlines an identified cluster that shows gradual increasing Z-score over time. (C) Two pie charts showing the proportional origin of differentially expressed proteins from panel A. Proteins already identified by MBR are represented by dark grey wedges, while exclusive identifications by DIA-ME are summed in blue wedges. The small pie displays the respective ratios for known IFN-γ-responsive proteins. (D) Heatmap of significantly up- and down-regulated proteins (p < 0.05, log_2_ fold change > 0.95) from panel A, showing their regulation over the course of treatment. Proteins indicated in blue were solely found after DIA-ME analysis. (E) Line plot of several selected proteins, taken from panel D, showing their gradual up-regulation over the course of treatment. (F) Gene set enrichment analysis of proteins that showed up-regulation after 24 hours (log_2_ fold change > 0.58) using MSigDB hallmarks. Bars on the right represent the degree of enrichment, while their colors specify the enrichment’s FDR. Bars on the left illustrate the size of the enriched term and are depicted for MBR (light grey) and DIA-ME (blue) analysis with numbers indicating the additional contribution of DIA-ME. Bold terms showed enrichment in 200-ng bulk analysis (see Fig. S7E).

In this way, DIA-ME allowed the additional identification of low-abundance proteins with high significance, thereby covering the induction of TAP2 and mTOR. Notably, the proteins whose expression could now be quantified over time (**Fig. 6D** and **E**), showed great similarity with our findings from 200-ng sample analysis (**Fig. S7H**). For instance, we observed gradual upregulation of TAP1/2, TAPBP and STAT1, which also grouped together with HLA-A and –C in a hierarchical cluster analysis (**Fig. 6B**). This cluster also contained calreticulin (CALR), a chaperone described to mediate the peptide loading process of TABP to the MHC class I complex (47) and the transmembrane protein CD40, which acts a mediator in the interaction of antigen-presenting cells with various immune cells (48). Furthermore, one of the strongest upregulated proteins was WARS1 (**Fig. 6A, B** and **E**), a tryptophanyl-tRNA synthetase that is known as a target of IFN-γ signaling (49, 50). We also recognized changes in the proteasome and found, as in the bulk data, the apparent restructuring of the 20S core particle by down-regulation of PSMB5 to PSMB7, and concomitant upregulation of their alternative subunits PMSB8 to PSMB10 that characterize the immunoproteasome (44–46) (**Fig. S8G**). Accordingly, overrepresentation analysis of upregulated proteins revealed the significant enrichment of several immuniologically relevant processes, including the signaling of IL6/JAK/STAT3, DNA repair and the underlying response to IFN-γ (**Fig. 6F**). Strikingly, DIA-ME analysis increased the size of enriched terms into an overall set that is highly similar to that obtained from the bulk data (cf. **Fig. 6F** and **S7E**). Collectively, these data demonstrate that DIA-ME not only improves proteome coverage in samples of exceedingly low input, but especially that it allows quantification of proteins that underlie the biological process at stake. Of note, these proteins were quantified from a 1,000-times lower input (200 pg) than the bulk experiment (200 ng), yet from a proteome whose coverage was only reduced by less than half (cf. **Fig. 5B** and **S7A**).

### Single-cell proteomic analysis by DIA-ME reveals protein co-expression and co-existence of cell states

Observing the benefit of DIA-ME for low-input proteomics (**Figure 6**), we next aimed to assess its performance in the analysis of actual single cells. We therefore used 24 hours IFN-γ-treated and control U-2 OS cells, and obtained a total number of 143 individual cells by FACS sorting. In addition, we collected instances of 10 cells to serve as ME samples during subsequent data analysis (**Table S7**).

Analogous to our observations from bulk samples, co-analysis with 10-cell samples via DIA-ME improved the proteomic depth from 496 to 575 median protein groups per individual cell (+16%) (**Fig. 7A**) and identified a total of 1,553 proteins (+41%) (**Fig. S9A**). Although proteome coverage was modest on an absolute scale, as sample preparation was conducted in-plate without further optimization, it fully served to demonstrate the benefit of DIA-ME. In particular, it also led to an increased number of identified peptides (**Fig. 7B**) and higher peptide coverage per protein (**Fig. 7C** and **D**), underpinning the advantage of this analysis for individual cells.

**Figure 7:**
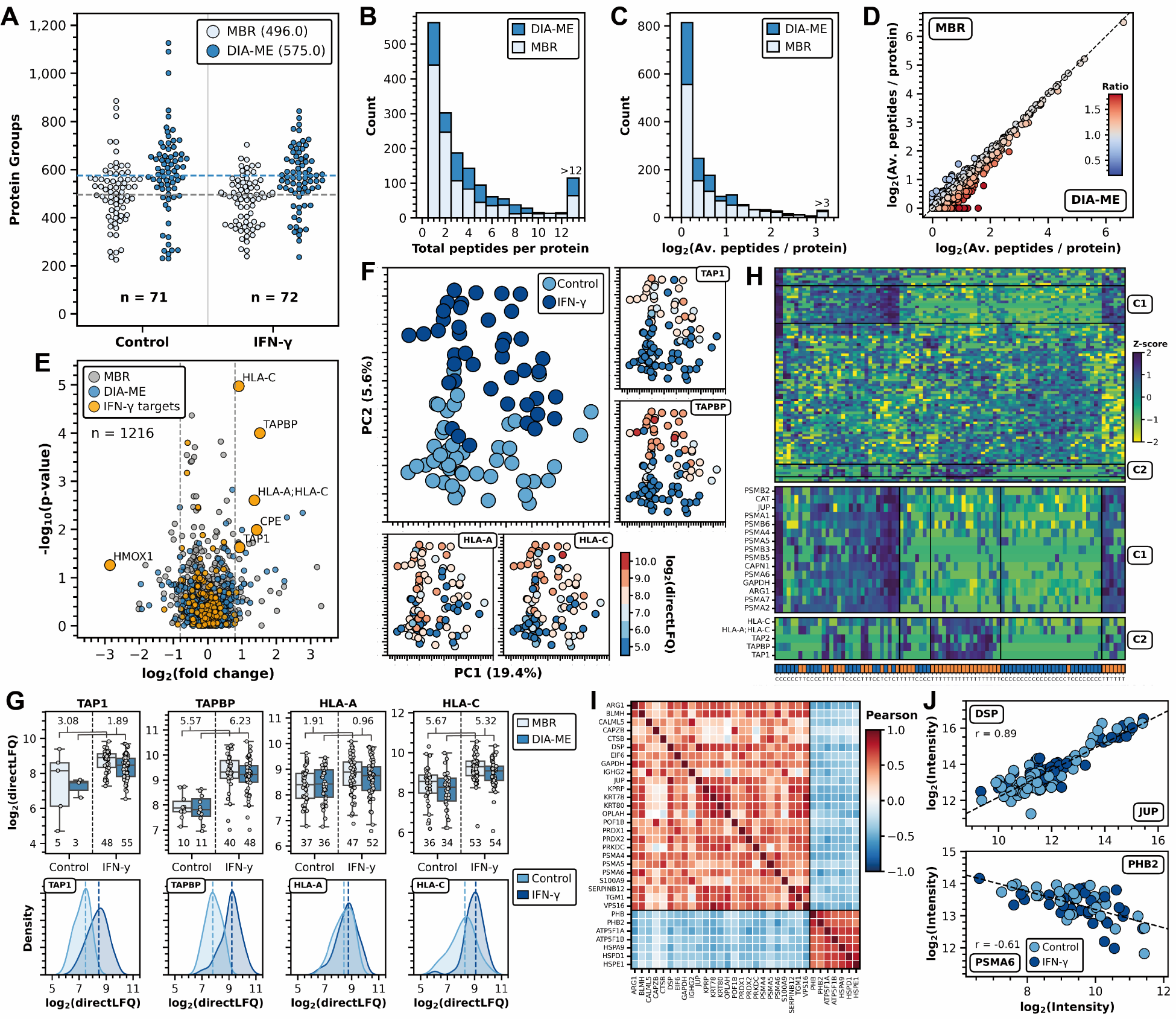
DIA-ME analysis of individual IFN-γ-treated U-2 OS cells reveals co-existence of metabolic states. (A) Number of identified protein groups per individual control cell and after 24 hours of IFN-γ treatment. Data was analyzed using conventional MBR (white) or DIA-ME (blue) using 10-cell MEs. The number of single cells per condition are indicated. (B) Histogram of total identified peptides per protein for both analyses (white: MBR; blue: DIA-ME). (C) Histogram of average number of peptides per protein (log2-transformed) for MBR (white) and DIA-ME analysis (blue). (D) Scatter plot of average peptide number per protein identified in MBR (y-axis) and DIA-ME analysis (x-axis). Colors of proteins represent the ratio of peptide identifications between the two analyses, showing higher sequence coverage in MBR (blue) or in DIA-ME (red). The black line indicates equal peptide numbers (ratio of 1). (E) Volcano analysis of Student’s t-test results between IFN-γ-treated and control cells. n: Number of identified differential expressions. Blue: differentially expressed proteins exclusively observed after DIA-ME analysis; yellow: protein known to be involved in the cellular response to IFN-γ. (F) Head-to-head evaluation of the four exemplary proteins (boxplots and kernel distributions) for MBR (white) and DIA-ME analysis (blue) and for control (light blue) and IFN-γ treatment (dark blue). Individual protein abundance per cell are shown as grey dots (number of cells indicated at the bottom). Numbers on top indicate p-values between populations in negative log_10_ scale. (G) PCA analysis of control and treated cells after DIA-ME analysis based known IFN-γ-responsive proteins. Detected intensities of four exemplary proteins are highlighted according to a color bar ranging from blue (low expression) to red (high expression). (H) Hierarchical cluster analysis by Euclidean distance of individual cells based on known IFN-γ-responsive proteins. Colors in the heatmap indicate quantitative changes compared to the protein’s median value across all cells by Z-score. Boxes indicate two clusters (C1, C2) that are enlarged at the bottom. Orange: IFN-γ-treated cell; blue: control cell. (I) Pearson correlation heatmap of highly correlated proteins among a minimum of 20 cells. (J) Examples of pairwise-correlated proteins JUP (x-axis) and DSP (y-axis) (top), and PSMA6 (x-axis) and PHB2 (y-axis) (bottom) as scatter plot reflecting respective expression levels in individual cells. Dark blue: IFN-γ-treated cell; blue: control cell. Linear regressions by Pearson are shown as dashed lines, the respective correlation r is indicated.

To investigate the proteome effect of IFN-γ and reduce the effect of missing values, we only retained data where at least 500 proteins per cell were identified after DIA-ME analysis (90 cells remained). Several of the IFN-γ-induced proteins observed in bulk and low-input samples were significantly upregulated in single-cell data as well (**Fig. 7E**). This comprised TAP1, TAPBP, HLA-A and HLA-C, which also drove the cluster separation between untreated and treated cells in PCA analysis (**Fig. 7F**). Accordingly, these proteins also displayed higher expression levels in treated cells, shown by a comparative analysis between both experimental settings. For instance, we observed a notable disparity in protein expressions within each condition, resulting in an almost 10-fold range of abundances among treated cells in case of TAP1 and TAPBP (**Fig. 7G**). Interestingly, protein levels in some of the cells were as low as in untreated cells, suggesting differences in their responsiveness, or that the attained expression levels depend on the initial level in the untreated cell. To discern distinct patterns of IFN-γ-responsive protein expression among cells, we conducted a hierarchical clustering analysis. This revealed two protein clusters, one of which comprised structural and catalytical subunits of the 20S proteasome (cluster C1), while the other consisted of proteins of the antigen presentation pathway, such as TAP1/2, TAPBP and HLA-C (cluster C2) (**Fig. 7H**). As expected from the previous analyses, cells in cluster C2 grouped together according to their experimental condition, showing upregulation of the respective proteins in treated cells. Remarkably, we observed that proteins in cluster C1 divided into two groups regardless of the experimental condition, including several proteasomal proteins and metabolic enzymes like catalase (CAT) and GAPDH (**Fig. 7H**). This indicates that cells with different metabolic activity co-exist in the initial cell population, yet such diversity does not dictate the responsiveness to IFN-γ. Likewise, it suggests that proteins of the proteasomal and enzymatic cluster are correlated in their expression, showing either collective up- or down-regulation.

To further explore the existence and degree of protein co-expression within the same cell, we performed a co-variation analysis across proteins that were identified in ≥ 20 cells, irrespective of the treatment with IFN-γ. This has the potential to provide insight into the correlation in function or regulation of expression that cannot be obtained from bulk populations and that is unique to single-cell analysis. Among 570 proteins, this revealed two clusters that showed high correlation within each cluster while showing inversed correlation between them (**Fig. S9B**), encompassing 111 proteins (**Fig. S9C** and **Table S8**). The most strongly co-varying proteins (**Fig. 7I**) include several complex-forming proteins, comprising the 20S proteasome subunit (PSMA4/5/6), the prohibitin complex (PHB1/2), ATP synthase F1 subunit (ATP5F1A/B), and the Hsp60-Hsp10 chaperonin complex (HSPD1/E1), but also functionally related proteins, e.g. the peroxiredoxins PRDX1/2, and the adherens junction proteins JUP and DSP (**Fig. 7J** and **S10**). Moreover, we observed strong anti-correlation between various protein pairs of diverse function (**Fig. 7I** and **J, Fig. S10**), suggesting mutually exclusive expression. Overrepresentation analysis of the two clusters revealed complementary cellular functions, showing the specific enrichment of degradative processes in cluster C1, such as the proteasome, DNA damage response and apoptosis (**Fig. S9D**), and proliferative processes in cluster C2, including glucose metabolism, TCA cycle, regulation of the mitotic cell cycle and formation of ATP (**Fig. S9E**). Given their inverse correlation, this suggests the co-existence of cells in two mutually exclusive metabolic states. Using network analysis, we further demonstrate that a hub of proteasome proteins drives the cluster harboring degradative processes, while the inversely related proteins that connect both clusters may provide interesting examples to infer novel functional relationships (**Fig. S9F**). For instance, we found mutually exclusive association between specific proteins involved in glycolysis, TCA cycle, oxidative stress and mitochondrial activity (**Fig. S9G**), while GAPDH (glyceraldehyde 3-phosphate dehydrogenase) appeared to be the hub protein in the network linking these respective terms (**Fig. S9F**). GAPDH is a crucial enzyme in glycolysis, however, has recently been described to divert glycolytic flux into the pentose phosphate pathway upon oxidation by intracellular hydrogen peroxide (H_2_O_2_) (51, 52), making the cells tolerant to oxidative stress (52). Our data may thus be explained by the interplay between these physiological processes including co-expression of GAPDH and PKM (pyruvate kinase M1/2) with oxidative response proteins, such as CAT and the peroxiredoxins PRDX1/2, resulting in reduced TCA and mitochondrial activity (**Fig. S9G**), and could be the consequence of detachment of cells, which is known to elevate endogenous oxidant levels (52–54). In a related context, we observed that the expression of cytosolic peroxiredoxin (PRDX)-1 and 2 anti-correlated with mitochondrial PRDX3 (**Fig. S9G**), showing different modes of expression regulation of these isoforms, potentially related to their complementary functionality in their respective sub-cellular locations. Other examples are serine proteases SERPINA1, -B3 and -B12 that negatively correlate with SERPINH1 (**Fig. S9F**), suggesting mutually exclusive expression to possibly regulate collagen formation. In addition, we found that proteasomal proteins negatively correlate with several dozens of other proteins (**Fig. 7I, Fig. S9F**), potentially indicating enzyme-substrate relationships. These and many other observations from co-expression analysis exemplify hypotheses enabled by single-cell proteome analysis, opening exciting novel avenues to investigate causality of these interactions.

In conclusion, we applied DIA-ME to successfully explore proteome dynamics in single cells upon IFN-γ treatment, using 10-cell samples as matching resource to improve proteome coverage. The concept is readily scalable to larger cell numbers, and is extendable to any other cell types and treatments to investigate proteome response and protein co-variation in other biological contexts.

## Discussion

In this study, we introduced DIA-ME as an experimental approach to increase proteome coverage and completeness in low-input and single-cell proteomics. DIA-ME employs a higher-input reference sample that is co-analyzed with low input samples of interest, and was designed to exploit the capabilities of DIA analysis tools such as DIA-NN and Spectronaut. In particular, we refer to the reference sample as matching enhancer because it serves to generate an augmented internal library in DIA analysis software for improved matching to features from low-input data, resulting in an increased number of protein identifications. This approach is easy to implement in existing proteomic pipelines as it just requires the analysis of a small set of higher input amounts along with the samples of interest. We determined that an ME sample containing 10x higher input (e.g. 10 cells along with single-cell samples) suffices to considerably increase proteome coverage and data completeness. In this regard, DIA-ME is advantageous over library-based DIA applications that require time-intensive generation of spectral libraries. Furthermore, DIA-ME makes economic use of potentially scarce biological material since only a few ME samples need to be analyzed along with any desired number of low input samples. Conceptually even one ME sample would be sufficient to introduce an adequate number of additional features to the search, however, slightly more replicates could be used to account for potential missing values, as we did in this study. In this sense, MEs of extreme time-points during IFN-γ treatment resulted in higher proteome coverage than MEs of individual time-points, with equivalent results when all time-points were used (**Fig. S8A**). This emphasizes that only a few MEs are required to cover the detectable peptide space, in theory along with a limitless number of single cell samples, making DIA-ME a highly scalable approach. This contrasts with the concept of carrier channels in TMT-experiments, where a sample at 20–100× the amount of a single-cell proteome should be added to every plex of 8 or 16 single cells. Moreover, the amount needs be carefully tuned to account for ratio compression and collection of sufficient number of ions to allow for appropriate peptide quantification (19). Thus, DIA-ME will be particularly advantageous in scenarios where cells of interest are scarce or hard to obtain, e.g. in clinical proteomics or developmental biology.

A crucial element of DIA-ME is its reliance on the feature matching functionality as implemented in DIA data analysis tools, and therefore we extensively evaluated the degree of false transfers, particularly in scenarios where low and high input samples are co-analyzed (**Fig. 3**). To this end, we used a two-proteome model and analyzed non-spiked and spiked samples together to assess the fidelity of the MBR function, as has been done previously for DDA in MaxQuant (55). This revealed that the quality both of protein identification and identity transfer was not affected in DIA-NN (FPR < 1% and FTR 1.0 – 1.8%, respectively), even when co-analyzing a 1-ng human sample with a 100-fold excess ME sample that contains 20% (i.e. 20 ng) *E.coli* peptides. Counterintuitively, FTRs even improved with increased size of the ME sample (**Fig. 3C**), which we hypothesize is the result of better spectral quality that enhances the matching process. Although we did not investigate this further, spectral data obtained from MEs may be much more similar to the data in low-input samples than to higher-input or even deep-fractionated libraries (56, 57), which has been reported to promote false discoveries due to the large query space (58, 59). Indeed, we observed that the average number of proteins does not increase with extensive database size beyond 10x DIA-ME (**Fig. 2B** and **Fig. 5B**). Collectively, these data show that the matching process in DIA-NN is resilient to the presence of highly abundant interfering peptides, while effectively discriminating false- and true-positive identifications. DIA-NN outperformed Spectronaut in both aspects, although this might be partly due to the lenient default filtering of the latter (**Fig. S5**). It will be interesting to identify the specific determinants in DIA-NN contributing to this observation, and to potentially improve its performance even further by tuning parameters especially for single-cell applications, e.g. with regard to the role of the TIMS dimension in diaPASEF data for the matching algorithm (27, 60). Until then, and as a practical implication, we found that 10x ME samples provide a good balance between increased proteome coverage and consistent identifications, even though the exact ratio may require further investigation.

With the notion that single-cell proteomics is still in its infancy, applications are diversifying from analytical studies that verify the ability to distinguish different cell types (18, 30, 32, 33, 61–64), to experiments that explore proteome changes induced by potent treatments such as LPS stimulation (35, 65) or that investigate more subtle effects e.g. during cell cycle (34) or in stem cell populations (31). Herein, we applied DIA-ME to investigate the proteome response to IFN- γ, which only induces a mild effect in U-2 OS cells as determined in a bulk experiment not limited by sample input (**Fig. S7**). When using low-input sample amounts either from diluted cell extracts (200 pg peptides) or from single cells, DIA-ME recapitulated the main observations of the bulk experiment by quantifying the up-regulation of known IFN- γ response proteins, showing very similar time-resolved expression profiles (cf. **Fig. S7H** and **6E**) and enriched gene sets (cf. **Fig. S7E** and **6F**). This indicates that DIA-ME can detect biologically relevant consequences of mild treatments in single cells. Of note, these results were reliant on proper normalization to ensure unaffected protein quantification, where the recently published directLFQ algorithm (40) showed the best quantitative accuracy among several tested normalization strategies (**Fig. S6**). The ability of DIA-ME to study biological processes at the single cell level is likely to be more pronounced with increased proteomic depth, e.g. when using sample preparation protocols optimized for single cells, and using more sensitive mass spectrometry technologies than those used here.

Finally, in our data we observed many examples of proteins whose expression was co- or anti-correlated within the same cell, indicating the co-existence of different cell states within the population of U2-OS cells (**Fig. 7I** and **J, S9B-G, S10**). The ability to obtain such patterns of co-varying proteins is a major perspective of single-cell proteomics to elucidate fundamental aspects that underlies heterogeneity within cell populations, to reveal regulatory processes in protein expression, or to infer other functional relationships between proteins (66). This concept has been successfully used in bulk proteomics to reveal novel functions of uncharacterized proteins (67), but it has greater potential when performed at the single-cell level, which does not suffer from masking effects due to heterogeneity of cell populations. Indeed, this has begun to be explored in recent single-cell studies, observing co-regulation of proteins that can be rationalized from their interaction in complexes (68) or their complementary involvement in energy metabolism (62). In our data, we observed many instances of proteins with inverse expression patterns, but with similar functions (e.g. in glycolysis, oxidative stress, proteases), suggesting mutually exclusive or complementary functionality. Although our data cannot infer causality of these relationships, they provide novel hypotheses that can be tested in future studies. From a systems perspective, it will be interesting to understand if modules of co- and anti-correlated proteins are maintained across different states of the same cell type, or across different cell types. DIA-ME should provide a powerful approach to conduct such studies in the future to increase our understanding of proteome regulation at the single-cell level.

## Material & Methods

### Preparation of *E.coli* peptides

Lyophilized *E.coli* K12 sample (Bio-Rad) was reconstituted in 50 mM triethylammonium bicarbonate (TEAB, pH 8.5, Sigma-Aldrich) buffer to a stock concentration of 1 µg/µL. 100 µg were incubated at 95 °C for 5 min and subsequently worked-up by SP3 (69): a mixture of 50:50 carboxylate-modified Sera-Mag SpeedBeads type A and B (Cytiva) were washed three times with ddH_2_O (Barstead GenPure, Thermo Scientific) on a magnetic rack and 1 mg of combined beads were added to the *E.coli* sample. For protein aggregation, the suspension was filled up with acetonitrile (ACN, Biosolve Chimie) to a concentration of 75% (v/v) and incubated at RT and 800 rpm for 20 min. Afterwards beads were rinsed twice with 1 mL 80% (v/v) ethanol (EtOH, VWR) and once with 800 µL ACN. During washing steps the suspension was homogenized to ensure adequate bead-solvent interaction. In the end, beads were air-dried for 2 min to get rid of ACN leftovers and resuspended in 30 µL digestion buffer (50 mM TEAB, pH 8.5 + 2 mM CaCl_2_ (Sigma-Aldrich)). Disulfide-bonds were reduced by 10 mM final concentration dithiothreitol (DTT, Biomol) and incubation at 37 °C and 600 rpm for 45 min. Next, bare cysteine residues were alkylated with 55 mM final chloroacetamide (CAA, Sigma-Aldrich) at RT and 600 rpm in the dark for 30 min. Finally, proteins were digested overnight at 37 °C and 800 rpm using sequencing-grade modified trypsin (Promega) at a protein-enzyme-ratio of 50:1. On the next day, the supernatant was transferred to a fresh tube and the beads were additionally washed with 50 µL 1% (v/v) trifluoroacetic acid (TFA, Biosolve Chimie) in ddH_2_O at RT and 800 rpm for 5 min to improve peptide recovery. The obtained peptide solution of pooled supernatant and washing fraction was subsequently cleaned-up on SepPak cartridges (Waters). The column was prepared according to the instructions of the manufacturer using 1 mL of ACN, solvent B (50% (v/v) ACN in ddH_2_O + 0.1% (v/v) formic acid (FA, Biosolve Chimie)) and solvent A (ddH_2_O + 0.1% (v/v) FA), respectively. Peptides were loaded on top of the column and washed twice with 1 mL solvent A before being eluted twice in 200 µL solvent B. Purified peptides were frozen at -80 °C, lyophilized in a freeze-dryer and afterwards reconstituted in solvent A. The resulting peptide concentration was determined in a colorimetric assay using bicichoninic acid (BCA) (Pierce, Thermo Scientific).

### Generation of two-species proteome model

Commercial HeLa S3 protein digest (Pierce, Thermo Scientific) was reconstituted in ddH_2_O + 0.1% (v/v) FA by gentle vortexing. For preparation of spiked human samples, defined amounts of the HeLa peptide standard were mixed with 5%, 10% and 20% (w/w) of prepared *E.coli* peptides to final concentrations of 0.5 ng/µL (n = 7), 2.5 ng/µL (n = 3), 5 ng/µL (n = 3) and 50 ng/µL (n = 3) per spike ratio. In addition, pure HeLa peptide standard was diluted in ddH_2_O + 0.1% (v/v) FA to a final concentration of 0.5 ng/µL (n = 7) (human-only).

### Cultivation of U-2 OS cells and Interferon-gamma treatment

Human osteosarcoma epithelial cells (U-2 OS cell line) were cultivated in DMEM high glucose medium (Gibco), supplemented with 2 mM L-glutamine, 10% (v/v) fetal bovine serum (Gibco), and an additional 2 mM GlutaMAX (Gibco). For the exploration of interferon gamma (IFN-γ) stimulation effects, U-2 OS cells underwent treatment with 50 ng/mL recombinant IFN-γ (Cell Signaling). The cytokine was diluted in 0.5% (w/v) bovine serum albumin (BSA, Serva).

In the context of preparing the time series experiment, cells were cultured in 10-cm dishes with three biological replicates, and IFN-γ treatments were administered when the cell confluence reached 70%. Cells were kept with the treatment for durations of 2 hours, 6 hours, 18 hours, and 24 hours. Additionally, distinct 0-hour and 24-hour time points were collected as controls.

After the respective incubation periods, the culture medium was carefully aspirated, and the cells underwent two rounds of washing with pre-warmed phosphate buffered saline (PBS, Sigma-Aldrich). Adherent cells were detached by gentle scraping in 5 ml of PBS, and subsequently centrifuged at 500 x g for 10 min. The resulting cell pellets were collected, rapidly frozen, and stored at -80 °C, preserving their integrity until their following preparation.

### Preparation of proteomic samples

For bulk analyses of 200-pg and 200-ng injections, U-2 OS cells obtained from cell culture were thawed and lysed in 50 mM TEAB, pH 8.5 (Sigma-Aldrich) buffer + 2% (v/v) Sodium dodecyl sulfate (SDS, Bio-Rad). Proteins were denatured at 95 °C and 600 rpm for 10 min before adding a final concentration of 2% (v/v) TFA (Biosolve Chimie) to the lysate. The reaction was quenched after 1 min by neutralizing pH with 3 M tris(hydroxymethyl)aminomethan (Tris, AppliChem) solution. Afterwards the lysate was sonicated for 15 cycles of 30 sec ON/OFF at 10 °C using a Bioruptor (Diagenode) and subsequently frozen at -80 °C. Prior to protein clean-up, the sample was thawed, centrifuged at 18,000 x g to separate remaining cell debris and transferred to a fresh tube. A total of 20 µg protein per sample (determined by BCA, Pierce Thermo Scientific) were employed in the SP3 protocol described above using 200 µg Sera-Mag SpeedBeads (Cytiva) and 200 µL of respective washing solutions. After digestion and subsequent bead washing, the acidified peptide solution was cleaned-up on self-packed StageTips (70) accommodating four discs of Empore C18 material (Merck Supelco). The material was prepared by consecutive application of pure ACN (Biosolve Chimie), solvent B (50% (v/v) ACN + 0.1% (v/v) FA (Biosolve Chimie) in ddH_2_O) and solvent A (0.1% (v/v) FA in ddH_2_O). Peptides were loaded on top of the column and washed twice with 1 mL solvent A, followed by elution in 100 µL solvent B. Purified peptides were frozen at -80 °C, lyophilized in a freeze-dryer and afterwards reconstituted in solvent A. The resulting peptide concentration was determined by BCA assay (Pierce, Thermo Scientific).

### FACS-sorting and preparation of individual cells

For our single-cell proteomics experiment, we utilized 24 hours control and IFN-γ-treated U-2 OS cells from the previously obtained pellets. Cells were prepared by gentle trypsin digestion (0.25%) to establish a homogeneous population of singularized cells. Approximately one million cells were diluted in 1.5 mL PBS (Sigma-Aldrich) and immediately administered to the cell sorting via a BD FACSAria III instrument (BD Biosciences), utilizing a 384-well plate configuration. Default settings for optical filters and mirrors were employed to facilitate the detection of the scattered signals. Parameters such as laser power, PMT-voltage settings, and gating were maintained at constant level throughput the entire experiment. Single cells were subjected to the sorting event using forward and side gating, while non-viable and dimerized cells were excluded. The sorting procedure was conducted at ambient room temperature. Sorted single and ten cells were directly collected in 2 µL of lysis buffer (50 mM TEAB, pH 8.5 + 0.025% (w/v) n-dodecyl-β-D-maltoside (DDM) (Sigma-Aldrich)) per well, immediately transferred to a chilled environment (ice-box), and subsequently prepared by the following in-plate protocol: cellular proteins were denatured at 70 °C and 600 rpm for 30 min, before adding 2 µL of 0.5 ng/µL trypsin (Promega) in digestion buffer (50 mM TEAB, pH 8.5) to the wells. Samples were incubated at 37 °C and 600 rpm for 1 h and afterwards acidified by 2 µL 0.5% (v/v) FA. The resulting volume of around 5 µL was transferred to a 96-well autosampler plate and subsequently injected into MS.

### Data acquisition (LC-MS/MS)

Desired peptide amounts of low-input samples were injected in 2 µL volume onto an analytical column (IonOpticks Aurora Series, 25 cm x 75 µm i.d. + CSI, 1.6 µm C18) using an EASY-nLC 1200 system (Thermo Scientific). Peptides were separated across a 15 min active gradient starting from 3.2% (v/v) ACN concentration in ddH_2_O (Biosolve Chimie) + 0.1% (v/v) FA to 13.6% (v/v) in 7.5 min to 20.0% (v/v) in 3.5 min to 28.0% (v/v) in 4 min at a flow rate of 300 nL/min and a temperature of 50 °C maintained by a column oven (Sonation). The LC system was connected to a timsTOF Pro mass spectrometer (Bruker Daltonics) via a nano-flow electrospray ionization (nano-ESI) source (Captive Spray, Bruker Daltonics). Analytes were ionized at 1,500 V capillary voltage, 3.0 L/min dry gas and 180 °C dry temperature. MS data was acquired in diaPASEF mode. In the TIMS, ions were accumulated to an IM constant 1/K_0_ of 1.7 V*s/cm^2^ and sequentially ramped from 1.3 to 0.75 V*s/cm^2^ over 100 ms in a locked duty cycle. Subsequent MS1 scans were performed from 200 to 1,700 m/z, while only precursors with a mass ratio of 475 to 1,000 m/z were considered for DIA window isolation. The range of 525 m/z was covered by equally sized windows of 25 Th width and 0.15 V*s/cm^2^ heights, which were combined to a total of 8 DIA scans (**Fig. S1D**) and resulted in 0.95 s cycle time. Precursor fragmentation was induced by IM-dependent collisional energies from 45 eV at 1/K_0_ of 1.3 V*s/cm^2^ to 27 eV at 0.75 V*s/cm^2^. The following ion detection was performed in high sensitivity mode for samples containing ≤ 50 ng peptide amount. In case of the analysis of single cells, data was acquired from an injection volume of 5 µL (entire sample) using the described method, but utilizing a flow rate of 200 nL/min.

The analysis of conventional bulk samples (200 ng) was performed on an Orbitrap Fusion Lumos Tribrid MS (Thermo Scientific) connected to an EASY-nLC 1200 system (Thermo Scientific). This LC system was configured in a trapping setup, comprising a Acclaim PepMap 100 trapping column (2 cm x 100 µm i.d., 5 µm 100 Å C18, Thermo Fisher Scientific) and a consecutive nanoEase M/Z Peptide BEH analytical column (25 cm x 75 µm i.d., 1.7 µm 130 Å C18, Waters). Peptides were loaded onto the trapping column at constant pressure of 800 bar, utilizing a total volume of 22 µL ddH_2_O + 0.1% (v/v) FA. Subsequently, peptides were separated in the analytical column along a 87 min gradient starting from 2.4% (v/v) ACN concentration in ddH_2_O + 0.1% (v/v) FA to 6.4% (v/v) in 4 min to 8.0% (v/v) in 2 min to 25.6% (v/v) in 68 min to 40.0% (v/v) in 12 min to 80% (v/v) in 1 min at a constant flow rate of 300 nL/min and a temperature of 45 °C maintained by a column oven (MonoSLEEVE, Analytical Sales and Services). Peptides were introduced into MS via a Nanospray flex ion source (Thermo Fisher Scientific) utilizing a Sharp Singularity nESI emitter (ID = 20 μm, OD = 365 μm, L = 7 cm, α = 7.5°, Fossiliontech) connected to a SIMPLE LINK UNO-32 (Fossiliontech). The emitter maintained a spray voltage of 2.5 kV, and the ion transfer tube capillary temperature was set to 275 °C. Data was acquired in DDA mode using MS1 full scans between 350 – 1500 m/z at a resolution of 120,000, and a maximum injection time (IT) of 32 ms with automatic gain control (AGC) of 3.0 x 10^6^. The top 20 most abundant precursors were isolated for fragmentations utilizing an isolation window of 2.0 m/z in the quadrupole, only allowing determined charges of 2 – 5. Dynamic exclusion was to set 40 sec with a mass tolerance of ±10 ppm. For MS2 acquisition, higher-energy collisional dissociation (HCD) was employed at 25% and the resulting fragments were acquired with an AGC target of 1.0 x 10^5^ or a maximum IT of 50 ms, covering a scan range from 200 – 2000 m/z in centroid mode.

### Raw data processing

After MS acquisition, DIA raw data was processed using library-free analysis in DIA-NN 1.8.0 or Spectronaut 18. We used a UniProtKB sequence table (fasta-file) of *Homo sapiens* (downloaded in September 2021, 20,386 reviewed proteins) for *in-silico* digestion and added an *Escherichia coli* K12 database (downloaded in April 2022, 4,529 reviewed proteins) in case of *E.coli*-spiked data.

For analyses in DIA-NN, we used deep learning-based prediction of spectra, retention times and ion mobilities and allowed a maximum of 2 missed cleavage sites (one for single-cell analysis) per peptide. We set peptide length range to 7 – 50 and precursor charge range to 2 – 4. Protein isoforms were grouped according to their protein names from fasta-files and we selected “any LC (high precision)” for precursor quantification. In case of the analysis of single cells, we moreover set the algorithm’s scan window to a value of 5 and MS1/Mass accuracies to 1.5e-05. For DIA-ME experiments, we combined raw files from low-input samples and MEs in the search and activated the match-between-runs (MBR) function. Analyses that are indicated as “MBR” did not contain ME samples, however, also have the MBR setting activated, whereas in analyses indicated as “Indiv”, raw files were searched individually without allowing feature matching. All other settings remained default, including data filtering at 1% precursor FDR.

For DIA-ME analyses in Spectronaut, we equivalently combined raw files from low-input samples and MEs in the search. Since Spectronaut matches features among samples by default, “MBR” refers to a search with unchanged settings without the addition of ME samples.

Analysis of DDA bulk data was performed in MaxQuant (v2.0.3.0) using a canonical *Homo sapiens* database (downloaded in November 2021, 20,394 reviewed proteins). Trypsin was specified as digestion enzyme, allowing up to 2 missed cleavage sites. Variable modifications included Oxidation (M), Acetyl (Protein N-term), and deamidation (NQ), while carbamidomethyl (C) was set as a fixed modification. For protein identification, the minimum unique peptides were set to 1, and peptide and protein hits were filtered at a 1% false discovery rate (FDR), with a minimum peptide length of 7 amino acids. The reversed sequences of the target database served as decoy for FDR calculation. The second peptide search option was activated. The MBR function was enabled with a matching time window of 0.4 min and an alignment time window set to 20 min. The “dependent peptides” function was deactivated. We activated label-free quantification and utilized unique and razor peptides for quantification. All other MaxQuant settings were maintained at their default configurations.

### Data analysis

Analysis of proteomic data was performed using Python code (version 3.9.12) in the Jupyter Notebook environment (v6.4.8) (71). To this end, we relied on the Pandas (v1.4.2) and NumPy (v1.21.6) packages for data handling, and used the Matplotlib (v3.5.1) and Seaborn (v0.11.2) packages for data plotting. Further statistical calculations, such as Student’s *t*-tests, hierarchical clustering and Gaussian approximation to model extracted ion chromatograms, were conducted by the SciPy (v1.7.3) package, while dimensional reduction and data imputation were performed using Scikit-learn (v1.0.2). Moreover, we plotted the UpSet plot using the UpSetPlot package (v0.6.1). For data normalization, we employed the directLFQ package (v0.2.8) (40) after excluding ME samples from the report table and filtering by Lib.Q.Value ≤ 0.01 and Lib.PG.Q.Value ≤ 0.01. Furthermore, to evaluate the performance of data normalization, we compared directLFQ to the DiaNN (v1.0.1) (26) and *iq* (v1.9.6) (39) packages using R Studio while applying the same filter criteria. Gene set enrichment analysis (GSEA) was conducted utilizing the R package clusterProfile (v3.12.0) (72) based on the “fgsea” algorithm. Data of ROC curves were computed in the Perseus platform (version 1.6.12) (73), calculating specificity and sensitivity while filtering on “PG.Q.Value” for DIA-NN or on “PG.QValue” for Spectronaut. Finally, illustrative figures of this work were created using BioRender.

For the assessment of erroneous feature assignment in DIA-NN and Spectronaut, we used the following calculations per replicate of non-spiked HeLa proteome (protein groups that shared human and *E.coli* proteins were excluded from the analysis):

#### False-positive rate

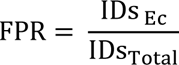

with *E.coli* protein identifications IDs _Ee_ and total protein identifications IDs _Total_.

#### False-discovery rate

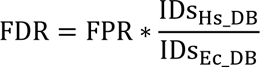

with *H.sapiens* and *E.coli* proteins present in the database (fasta) IDs _Hs_DB_ and IDs_Ee_DB_.

#### False-transfer rate

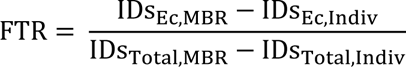

with *E.coli* protein identifications with and without activation of “MBR” IDs_Ee,MBR_ and IDs_Ee,Indiv_, and total protein identifications with and without activation of “MBR” IDs_Total,MBR_ and IDs_Total,Indiv_.

#### Species-specific transfer rate

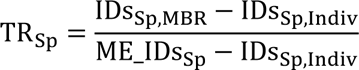

with species-specific protein identifications with and without activation of “MBR” IDs_Sp,MBR_ and IDs_Sp,Indiv_, and species-specific protein identifications in the matching enhancer source (ME) (after analyzing it without the target dataset) ME_IDs_Sp_.

For the similarity evaluation of DIA-ME and MBR data sets, we used following calculations:

#### Overlap (Szymkiewicz-Simpson) coefficient

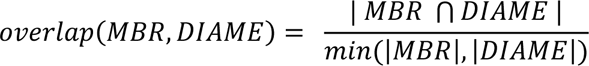

#### Jaccard similarity coefficient

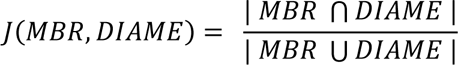

For the single-cell dataset, we performed a co-variation analysis by calculating the Pearson correlation between all identified proteins, tolerating a minimum of 20 pairwise observations (**Table S8**), and clustering the resulting dataset based on Euclidean distance. We identified two clusters and subsequently created a protein-protein interaction network using the Cytoscape software (v3.9.1) (74), leveraging the KEGG database for pathway information and the ClueGO plug-in for functional annotation. The interaction network was constructed by importing the protein list of both co-expression clusters and enriching it with relevant pathway information utilizing the KEGG database. Further, we employed the ClueGO plug-in to functionally annotate and highlight enriched pathways within the network. To ensure robustness of the enrichment procedures, a significance threshold (p-value ≤ 0.01) was applied. The resulting network was directly visualized in Cytoscape.

### Data annotation, filtering and imputation

For the analysis of IFN-γ-treatment experiments (dilution and single-cells), we filtered the protein intensity table on a contaminants list that we extracted from MaxQuant (v2.0.1). Afterwards we annotated proteins using a database that comprised known IFN-γ-responsive proteins. This database was retrieved from STRING database (https://string-db.org/) using IFN-γ-related signaling keywords, and extracting protein-protein interactions with confidence > 0.7 interaction score (75). Simultaneously, the Gene Ontology Consortium (http://geneontology.org/) provided annotations for relevant molecular functions and biological processes related to IFN-γ signaling. The extracted data from both sources were integrated based on common identifiers, and proteins identified in both databases were cross-referenced to ensure consistency (**Table S6**).

For the 200-pg dilution experiment, the obtained protein data was filtered on proteins that were identified at least three times per time-point and imputed the remaining missing values using k-nearest neighbors algorithm before conducting hierarchical clustering. Furthermore, principal component analysis was performed solely on proteins that showed full data completeness.

Following the DIA-ME analysis of single-cells, we determined the number of protein groups per individual cell and filtered for those cells that showed more than 500 identified proteins. The remaining data was used for the differential expression and co-variation analysis. Moreover, we excluded proteins that were identified in less than 30% of cells for principal component analysis and hierarchical clustering of known IFN-γ-responsive proteins and imputed missing values by the lowest intensity value present in the dataset.

### Data availability

Acquired raw LC-MS/MS files and report files of conducted data searches were deposited to the ProteomeXchange Consortium via the PRIDE partner repository with the dataset identifiers PXD048178 (*E.coli*-spiked data), PXD048162 (bulk IFN-γ experiment), PXD048175 (low-input IFN-γ experiment) and PXD048179 (single-cell data).

## Acknowledgement

This work was funded in part by the Federal Ministry of Education and Research (BMBF) and the Ministry of Science Baden-Württemberg within the framework of the Excellence Strategy of the Federal and State Governments of Germany. We specially thank the scOpenLab of the German Cancer Research Center (DKFZ) headed by Jan-Philipp Mallm for their help with FACS sorting and printing of single cells.

## References

1. Elowitz MB, Levine AJ, Siggia ED, Swain PS. Stochastic Gene Expression in a Single Cell. Science. 2002;297:1183–6.

2. Chen X, Teichmann SA, Meyer KB. From Tissues to Cell Types and Back: Single-Cell Gene Expression Analysis of Tissue Architecture. Annu Rev Biomed Data Sci. 2018;1:29–51.

3. Larsson AJM, Johnsson P, Hagemann-Jensen M, Hartmanis L, Faridani OR, Reinius B, et al. Genomic encoding of transcriptional burst kinetics. Nature. 2019;565:251–4.

4. Rodriguez J, Ren G, Day CR, Zhao K, Chow CC, Larson DR. Intrinsic Dynamics of a Human Gene Reveal the Basis of Expression Heterogeneity. Cell. 2019;176:213–26.

5. Raj A, van Oudenaarden A. Nature, Nurture, or Chance: Stochastic Gene Expression and Its Consequences. Cell. 2008;135:216–26.

6. Crow M, Paul A, Ballouz S, Huang ZJ, Gillis J. Characterizing the replicability of cell types defined by single cell RNA-sequencing data using MetaNeighbor. Nat Commun. 2018;9:884.

7. Haas S, Trumpp A, Milsom MD. Causes and Consequences of Hematopoietic Stem Cell Heterogeneity. Cell Stem Cell. 2018;22(5):627–38.

8. Franks A, Airoldi E, Slavov N. Post-transcriptional regulation across human tissues. PLoS Comp Biol. 2017;13(5):e1005535.

9. Buccitelli C, Selbach M. mRNAs, proteins and the emerging principles of gene expression control. Nat Rev Genet. 2020;21:630–44.

10. Levy E, Slavov N. Single cell protein analysis for systems biology. Essays Biochem. 2018;62(4):595–605.

11. Kelly RT. Single-cell Proteomics: Progress and Prospects. Mol Cell Proteomics. 2020;19(11):1739–48.

12. Bennett HM, Stephenson W, Rose CM, Darmanis S. Single-cell proteomics enabled by next-generation sequencing of mass spectrometry. Nat Methods. 2023;20:363–74.

13. Zhu Y, Piehowski PD, Zhao R, Chen J, Shen Y, Moore RJ, et al. Nanodroplet processing platform for deep and quantitative proteome profiling of 10–100 mammalian cells. Nat Commun. 2018;9:882.

14. Gebreyesus ST, Siyal AA, Kitata RB, Chen ES-W, Enkhbayar B, Angata T, et al. Streamlined single-cell proteomics by an integrated microfluidic chip and data-independent acquisition mass spectrometry. Nat Commun. 2022;13:37.

15. Masuda T, Inamori Y, Furukawa A, Yamahiro M, Momosaki K, Chang C-H, et al. Water Droplet-in-Oil Digestion Method for Single-Cell Proteomics. Anal Chem. 2022;94:10329–36.

16. Matsumoto C, Shao X, Bogosavljevic M, Chen L, Gao Y. Automated container-less cell precessing method for single-cell proteomics. bioRxiv preprint. 2022.

17. Petelski AA, Emmott E, Leduc A, Huffman RG, Specht H, Perlman DH, et al. Multiplexed single-cell proteomics using SCoPE2. Nat Protoc. 2021;16:5398–425.

18. Budnik B, Levy E, Harmange G, Slavov N. SCoPE-MS: mass spectrometry of single mammalian cells quantifies proteome heterogeneity during cell differentiation. Genome Biol. 2018;19:161.

19. Cheung TK, Lee C-Y, Bayer FP, McCoy A, Kuster B, Rose CM. Defining the carrier proteome limit for single-cell proteomics. Nat Methods. 2021;18:76–83.

20. Ctortecka C, Stejskal K, Krššáková G, Mendjan S, Mechtler K. Quantitative Accuracy and Precision in Multiplexed Single-Cell Proteomics. Anal Chem. 2022;94:2434–43.

21. Ye Z, Batth TS, Rüther P, Olsen JV. A deeper look at carrier proteome effects for single-cell proteomics. Commun Biol. 2022;5:150.

22. Ludwig C, Gillet L, Rosenberger G, Amon S, Collins BC, Aebersold R. Data-independent acquisition-based SWATH-MS for quantitative proteomics: a tutorial. Mol Syst Biol. 2018;14:e8126.

23. Meier F, Brunner A-D, Frank M, Ha A, Bludau I, Voytik E, et al. diaPASEF: parallel accumulatiom- serial fragmentation combined with data-independent acquisition. Nat Methods. 2020;17:1229–36.

24. Gillet LC, Navarro P, Tate S, Röst H, Selevsek N, Reiter L, et al. Targeted Data Extraction of the MS/MS Spectra Generated by Data-independent Acquisition: A New Concept for Consistent and Accurate Proteome Analysis. Mol Cell Proteomics. 2012;11:O111.016717.

25. Bruderer R, Bernhardt OM, Gandhi T, Miladinović SM, Cheng L-Y, Messner S, et al. Extending the Limits of Quantitative Proteome Profiling with Data-Independet Acquisition and Application to Acetaminophen-Treated Three-Dimensional Liver Microtissues. Mol Cell Proteomics. 2015;14(5):1400–10.

26. Demichev V, Messner CB, Vernardis SI, Lilley KS, Ralser M. DIA-NN: neural networks and interference correction enable deep proteome coverage in high throughput. Nat Methods. 2020;17:41–4.

27. Demichev V, Szyrwiel L, Yu F, Teo GC, Rosenberger G, Niewienda A, et al. dia-PASEF data analysis using FragPipe and DIA-NN for deep proteomics of low sample amounts. Nat Commun. 2022;13:3944.

28. Tsou C-C, Avtonomov D, Larsen B, Tucholska M, Choi H, Gingras A-C, et al. DIA-Umpire: comprehensive computational framework for data-independent acquisition proteomics. Nat Methods. 2015;12(3):258–71.

29. 29. Biognosys. Spectronaut 15 User Manual 2021 [Available from: http://files.biognosys.ch/058_Spectronaut/ReleaseMaterial/00_Manual/Spectronaut15_UserManual.pdf.

30. Webber KGI, Truong T, Johnston SM, Zapata SE, Liang Y, Davis JM, et al. Label-Free Profiling of up to 200 Single-Cell Proteomes per Day Using a Dual-Column Nanoflow Liquid Chromatography Platform. Anal Chem. 2022;94:6017–25.

31. Petrosius V, Aragon-Fernandez P, Üresin N, Kovacs G, Phlairaharn T, Furtwängler B, et al. Exploration of cell state heterogeneity using single-cell proteomics through sensitivity-tailored data-independent acquisition. Nat Commun. 2023;14:5910.

32. Wang Y, Lih T-SM, Chen L, Xu Y, Kuczler MD, Cao L, et al. Optimized data-independent acquisition approach for proteomic analysis at single-cell level. Clin Proteom. 2022;19:24.

33. Matzinger M, Müller E, Dürnberger G, Pichler P, Mechtler K. Robust and Easy-to-Use One-Pot Workflow for Label-Free Single-Cell Proteomics. Anal Chem. 2023;95:4435–45.

34. Brunner A-D, Thielert M, Vasilopoulou C, Ammar C, Coscia F, Mund A, et al. Ultra-high sensitivity mass spectrometry quantifies single-cell proteome changes upon pertubation. Mol Syst Biol. 2022;18(3):e10798.

35. Woo J, Clair GC, Williams SM, Feng S, Tsai C-F, Moore RJ, et al. Three-dimensional feature matching improves coverage for single-cell proteomics based on ion mobility filtering. Cell Syst. 2022;13(5):426–34.

36. Derks J, Leduc A, Wallmann G, Huffman RG, Willetts M, Khan S, et al. Increasing the throughput of sensitive proteomics by plexDIA. Nat Biotechnol. 2023;41:50–9.

37. Li Y, Li H, Xie Y, Chen S, Qin R, Dong H, et al. An Integrated Strategy for Mass Spectrometry-Based Multiomics Analysis of Single Cells. Anal Chem. 2021;93:14059–67.

38. Senavirathna L, Ma C, Chen R, Pan S. Spectral Library-Based Single-Cell Proteomics Resolves Cellular Heterogeneity. Cells. 2022;11:2450.

39. Pham TV, Henneman AA, Jimenez CR. iq: an R package to estimate relative protein abundances from ion quantification in DIA-MS-based proteomics. Bioinformatics. 2020;36(8):2611–3.

40. Ammar C, Schessner JP, Willems S, Michaelis AC, Mann M. Accurate Label-Free Quantification by directLFQ to Compare Unlimited Numbers of Proteomes. Mol Cell Proteomics. 2023;22(7):100581.

41. Schroder K, Hertzog PJ, Ravasi T, Hume DA. Interferon-γ: an overview of signals, mechanisms and functions. J Leukocyte Biol. 2004;55(2):163–89.

42. Platanias LC. Mechanisms of type-I and type-II-interferon-mediated signalling. Nat Rev Immunol. 2005;5:375–86.

43. Kak G, Raza M, Tiwari BK. Interferon-gamma (IFN-γ): Exploring its implications in infectious diseases. Biomol Concepts. 2018;9:64–79.

44. Rock KL, Goldberg AL. Degradation of cell proteins and the generation of MHC class I-presented peptides. Annu Rev Immunol. 1999;17:739–79.

45. Früh K, Yang Y. Antigen presentation by MHC class I and its regulation by interferon γ. Curr Opin Immunol. 1999;11:76–81.

46. Nandi D, Tahiliani P, Kumar A, Chandu D. The ubiquitin-proteasome system. J Biosci (Bangalore). 2006;31(1):137–55.

47. Williams A, Peh C, Elliott T. The cell biology of MHC class I antigen presentation. Tissue Antigens. 2002;59(1):3–17.

48. Elgueta R, Benson MJ, de Vries VC, Wasiuk A, Guo Y, Noelle RJ. Molecular mechanism and function of CD40/CD40L engagement in the immune system. Immunol Rev. 2009;229(1):152–72.

49. Biros E, Moran CS. Mini tryptophanyl-tRNA synthetase is required for a synthetic phenotype in vascular smooth muscle cells induced by IFN-γ-mediated β2-adrenoceptor signaling. Cytokine. 2020;127:154940.

50. Biros E, Vangaveti V, Moran CS. Mini-TrpRS is essential for IFNγ-induced monocyte-derived giant cell formation. Cytokine. 2021;142:155486.

51. van der Reest J, Lilla S, Zheng L, Zanivan S, Gottlieb E. Proteome-wide analysis of cysteine oxidation reveals metabolic sensitivity to redox stress. Nat Commun. 2018;9:1581.

52. Talwar D, Miller CG, Grossmann J, Szyrwiel L, Schwecke T, Demichev V, et al. The GAPDH redox switch safeguards reductive capacity and enables survival of stressed tumour cells. Nat Metab. 2023;5:660–76.

53. Schafer ZT, Grassian AR, Song L, Jiang Z, Gerhart-Hines Z, Irie HY, et al. Antioxidant and oncogene rescue of metabolic defects caused by loss of matrix attachment. Nature. 2009;461:109–13.

54. Jiang L, Shestov AA, Swain P, Yang C, Parker SJ, Wang QA, et al. Reductive carboxylation supports redox homeostasis during anchorage-independent growth. Nature. 2016;532:255–8.

55. Lim MY, Paulo JoA, Gygi SP. Evaluating False Transfer Rates from the Match-between-Runs Algorithm with a Two-Proteome Model. J Proteome Res. 2019;18:4020–6.

56. Zhong C-Q, Wu R, Chen X, Wu S, Shuai J, Han J. Systematic Assessment of the Effect of Internal Library in Targeted Analysis of SWATH-MS. Anal Chem. 2020;19:477–92.

57. Boekweg H, Van der Watt D, Truong T, Johnston SM, Guise AJ, Plowey ED, et al. Features of Peptide Fragmentation Spectra in Single-Cell Proteomics. J Proteome Res. 2022;21:182–8.

58. Rosenberger G, Bludau I, Schmitt U, Heuesel M, Hunter CL, Liu Y, et al. Statistical control of pepitde and protein error rates in large-scale targeted data-independent acquisition analyses. Nat Methods. 2017;14(9):921–9.

59. Siyal AA, Chen ES-W, Chan H-J, Kitata RB, Yang J-C, Tu H-L, et al. Sample Size-Comparable Spectral Library Enhances Data-Independent Acquisition-Based Proteome Coverage of Low-Input Cells. Anal Chem. 2021;93:17003–11.

60. Prianichnikov N, Koch H, Koch S, Lubeck M, Heilig R, Brehmer S, et al. MaxQuant Software for Ion Mobility Enhanced Shotgun Proteomics. Mol Cell Proteomics. 2020;19(6):1058–69.

61. Liang Y, Acor H, McCown MA, Nwosu AJ, Boekweg H, Axtell NB, et al. Fully Automated Sample Processing and Analysis Workflow for Low-Input Proteome Profiling. Anal Chem. 2021;93:1658–66.

62. Leduc A, Huffman RG, Cantlon J, Khan S, Slavov N. Exploring functional protein covariation across single cells using nPOP. Genome Biol. 2022;23:261.

63. Ctortecka C, Hartlmayr D, Seth A, Mendjan S, Tourniaire G, Udeshi ND, et al. An Automated Nanowell-Array Workflow for Quantitative Multiplexed Single-Cell Proteomics Sample Preparation at High Sensitivity. Mol Cell Proteomics. 2023;22(12):100665.

64. Williams SM, Liyu AV, Tsai C-F, Moore RJ, Orton DJ, Chrisler WB, et al. Automated Coupling of Nanodroplet Sample Preparation with Liquid Chromatography−Mass Spectrometry for High-Throughput Single-Cell Proteomics. Anal Chem. 2020;92:10588–95.

65. Huffman RG, Leduc A, Wichmann C, Giola MD, Borriello F, Specht H, et al. Prioritized mass spectrometry increases the depth, sensitivity and data completeness of single-cell proteomic. Nat Methods. 2023;20:714–22.

66. Slavov N. Learning from natural variation across the proteomes of single cells. PLoS Biol. 2022;20(1):e3001512.

67. Kustatscher G, Grabowski P, Schrader TA, Passmore JB, Schrader M, Rappsilber J. Co-regulation map of the human proteome enables identification of protein functions. Nat Biotechnol. 2019;37:1361–71.

68. Hu M, Zhang Y, Yuan Y, Ma W, Zheng Y, Gu Q, et al. Correlated Protein Modules Revealing Functional Coordination of Interacting Proteins Are Detected by Single-Cell Proteomics. J Phys Chem B. 2023;127:6006–14.

69. Hughes CS, Moggridge S, Müller T, Sorensen PH, Morin GB, Krijgsveld J. Single-Pot, solid-phase-enhanced sample preparation for proteomic experiments. Nat Protoc. 2019;14:68–85.

70. Rappsilber J, Mann M, Ishihama Y. Protocol for micro-purification, enrichment, pre-fractionation and storage of peptides for proteomics using StageTips. Nat Protoc. 2007;2(8):1896–906.

71. Kluyver T, Ragan-Kelley B, Pérez F, Granger B, Bussonnier M, Frederic J, et al. Jupyter Notebooks - a publishing format for reproducible computational workflows. In: Loizides F, Schmidt B, editors. Positioning and Power in Acadamic Publishing: Players, Agents and Agendas: IOS Press; 2016. p. 87–90.

72. Yu G, Wang L-G, Han Y, He Q-Y. clusterProfiler: an R Package for Comparing Biological Themes Among Gene Clusters. OMICS: J Integrative Biol. 2012;16:284–7.

73. Tyanova S, Temu T, Sinitcyn P, Carlson A, Hein MY, Geiger T, et al. The Perseus Computation Platform for Comprehensive Analysis of (Prote)omics Data. Nat Methods. 2016;13(9):731–40.

74. Shannon P, Markiel A, Ozier O, Baliga NS, Wang JT, Ramage D, et al. Cytoscape: A Software Environment for Integrated Models of Biomolecular Interaction Networks. Genome Res. 2003;13(11):2498–504.

75. Szklarczyk D, Gable AL, Lyon D, Junge A, Wyder S, Huerta-Cepas J, et al. STRING v11: protein– protein association networks with increased coverage, supporting functional discovery in genome-wide experimental datasets. Nucleic Acids Res. 2019;47:607–13.

