## Supplementary Figures for "Enhanced feature matching in single-cell proteomics characterizes response to IFN-γ and reveals co-existence of different cell states"

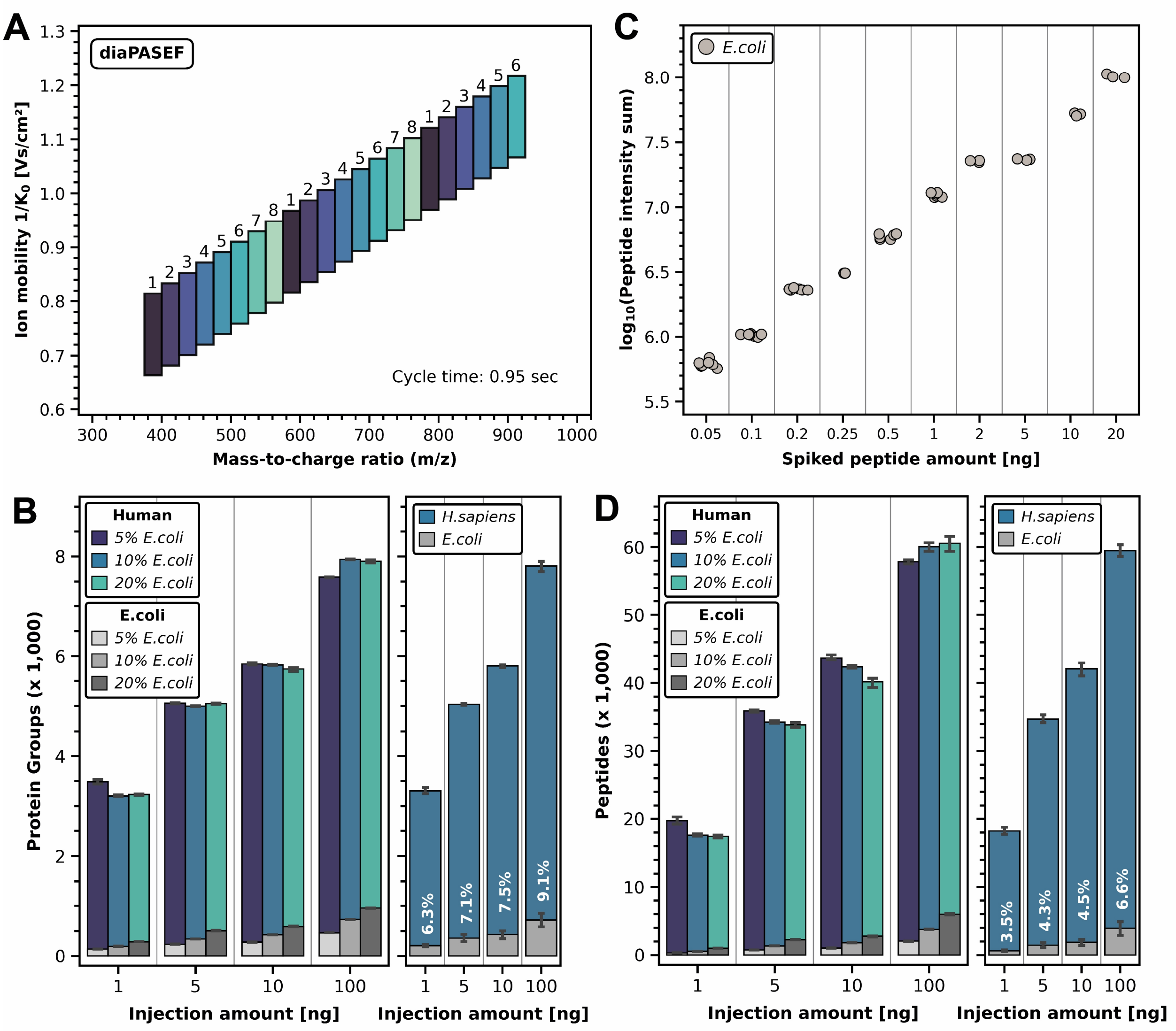


**Figure S1: Measurement and preparation of ME samples.** (A) Arrangement of diaPASEF windows for the measurement of all samples in this work. In MS1, the method covers an m/z range of 550 Th, ranging from 375 m/z to 925 m/z, and an ion mobility range of about 0.56 $1/K_0$, ranging from 1.22 to 0.66. Windows with equal numbers were combined for precursor fragmentation. (B) Average number of identified human and *E.coli* protein groups in the spiked ME samples. Left: Protein groups per injection amount (1 – 100 ng) and *E.coli* spiking ratio (5%, 10% and 20%), respectively. Right: Average protein groups per injection amount (1 – 100 ng) among spiking ratios. Percentages of *E.coli* proteins among average protein group numbers are indicated. (C) Cumulative peptide intensities ($log_{10}$-transformed) per spiked peptide amount, i.e. spiking ratio multiplied with the respective injection amount. (D) Same as (B), but for identified peptides.

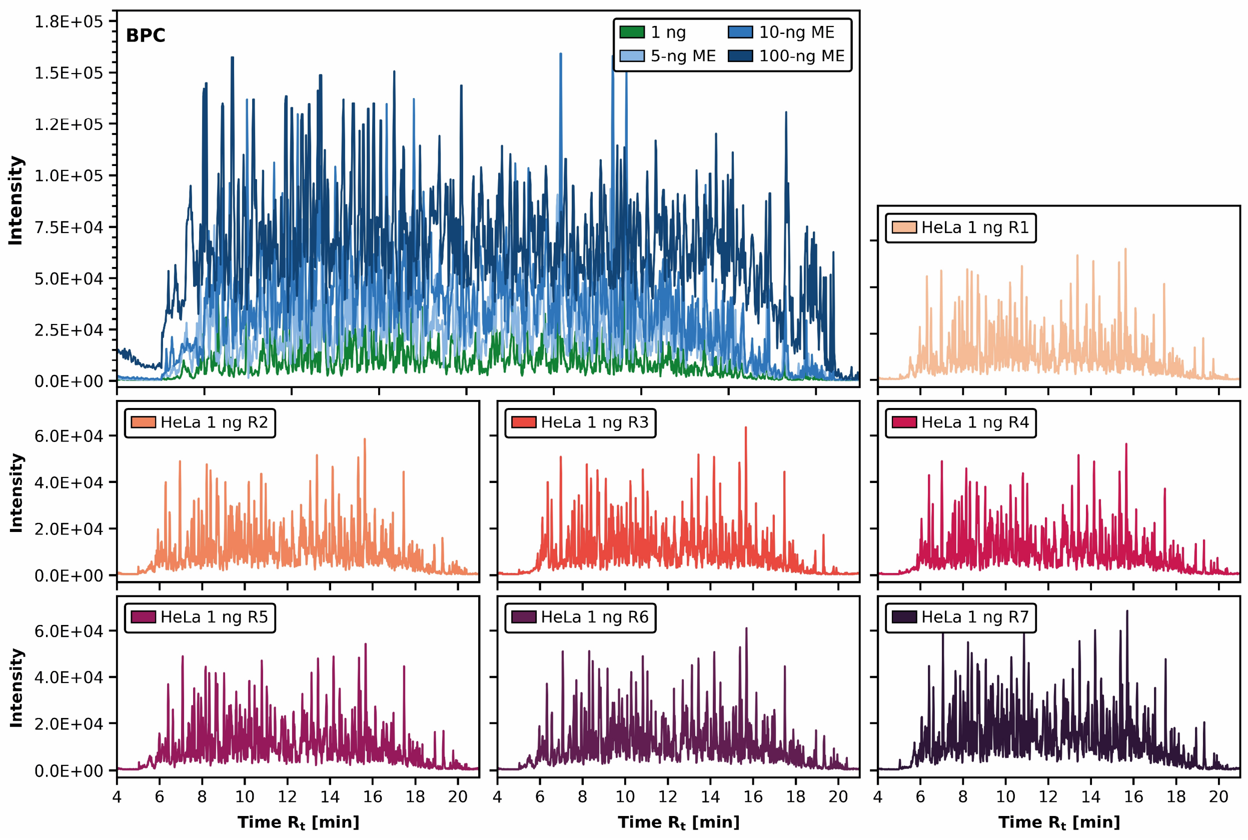


**Figure S2: Base peak chromatogram (BPC) of low-input HeLa replicates and MEs.** Top panel: overlay of exemplified BPCs of one spiked 5-ng, 10-ng and 100-ng replicate (MEs), respectively, and one non-spiked 1-ng replicate. Bottom panels: BPCs of all seven non-spiked 1-ng replicates.

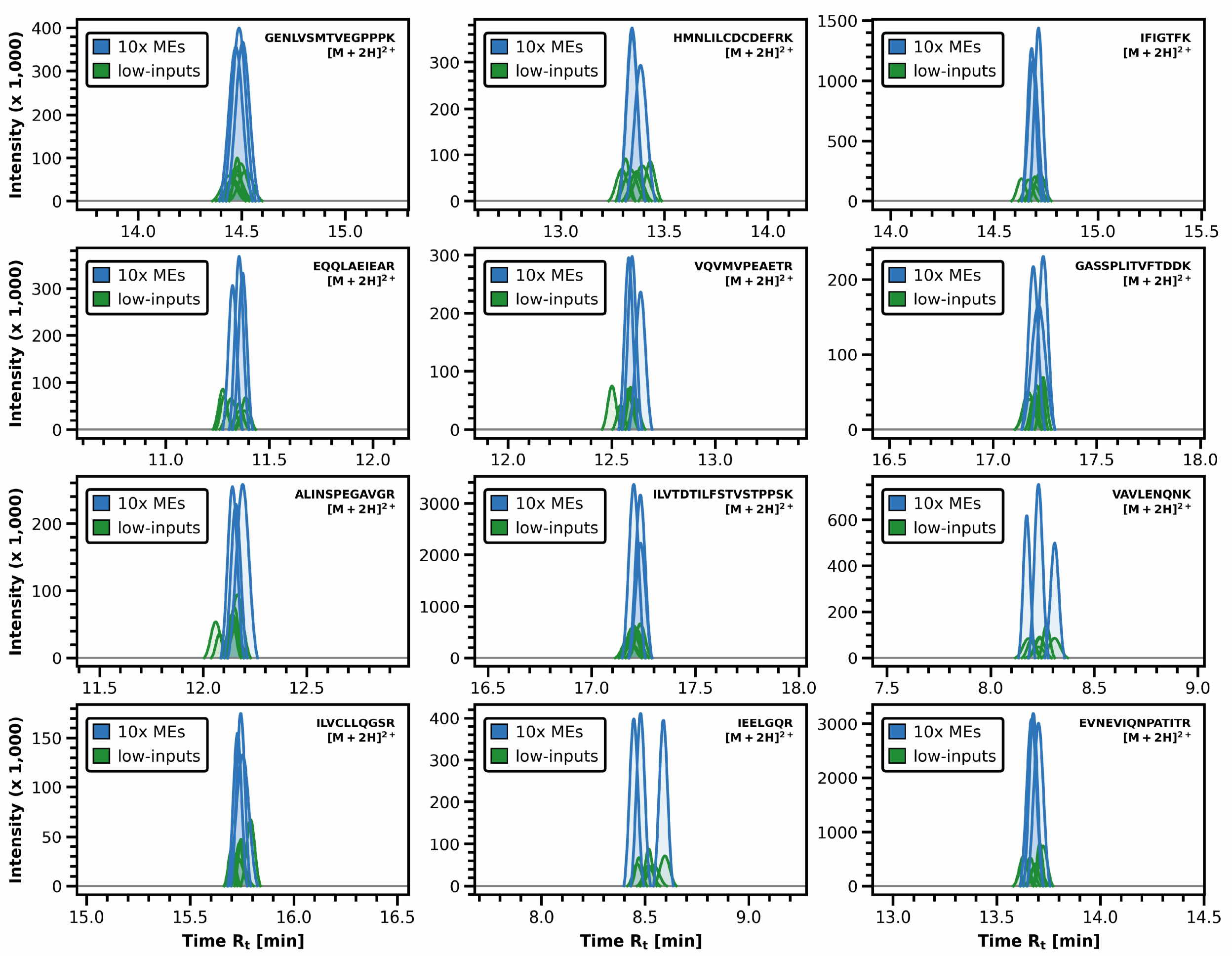


**Figure S3: Superimposed elution peaks of several peptides identified by DIA-ME analysis.** Peaks are shown for peptides of different proteins that were identified in the 10-ng ME samples (i.e. 10x ME, blue) and were matched to the low-input replicates (1 ng, green), resulting in full data completeness. Peptide sequences and their respective charge state are indicated. Elution profiles were calculated by Gaussian approximation.

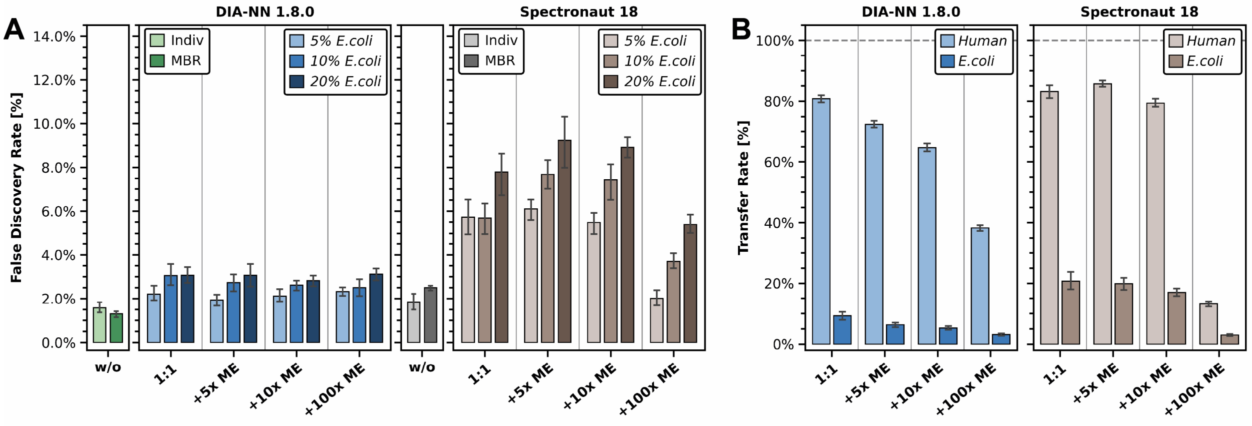


**Figure S4: False discovery and species-specific transfer rate in non-spiked 1-ng samples.** (A) Estimated protein false discovery rate, i.e. percentage of detected *E.coli* proteins corrected by the database sizes (see methods), in non-spiked 1-ng HeLa samples for different types of data analysis and DIA software. Analysis without spiked samples are indicated in green and grey (light-: without MBR, dark-: MBR), while analyses together with spiked samples are indicated in blue (DIA-NN) and brown (Spectronaut). The shade of the color indicates the *E.coli* spiking ratio. Error bars are shown as mean ± sd. (B) Species-specific transfer rate of human (light) and *E.coli* (dark) proteins from spiked samples to non-spiked samples in different analyses for DIA-NN (blue) and Spectronaut (brown). The transfer rate describes that percentage/probability of a protein to be transferred from a donor to an acceptor sample, if it was not identified in the latter before. Error bars are shown as mean ± sd.

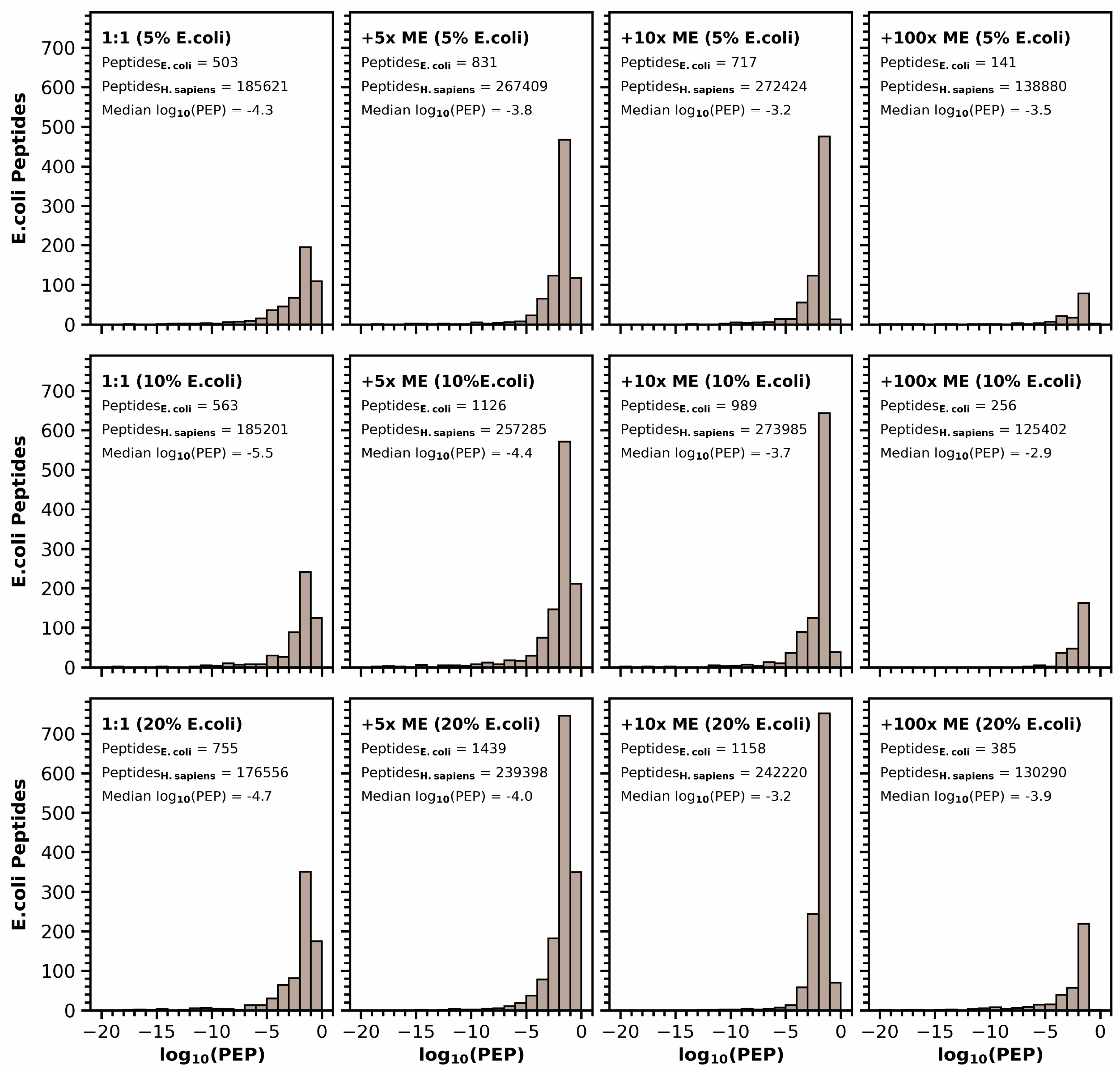


**Figure S5: PEP filtering in Spectronaut for low-input DIA data.** Histograms of identified *E.coli* peptides among non-spiked HeLa replicates along their posterior error probability (PEP) scores ($log_{10}$-transformed) after data co-analysis. Numbers of cumulative *E.coli* and *H.sapiens* peptides, as well as the observed median PEP scores per analysis are indicated. Note that the default filter for peptides in Spectronaut are PEP values of ≤ 0.1, thus requiring more stringent filtering for low-input applications.

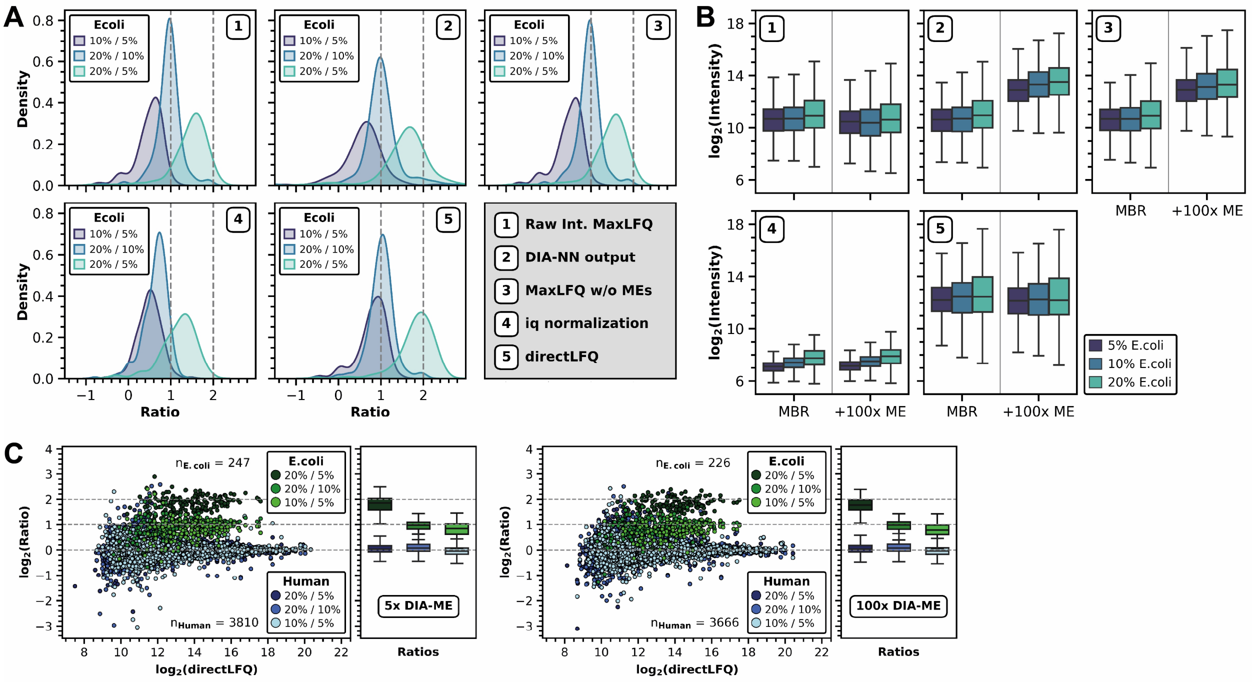


**Figure S6: Comparison of different normalization strategies for low-input DIA data after DIA-NN search.** (A) Mean *E.coli* protein quantity ratios between different spiking amounts after MBR analysis (without MEs) for different data normalizations. 1: DiaNN *R* package using raw peptide intensities; 2: protein output table of DIA-NN; 3: DiaNN *R* package using pre-normalized peptide intensities; 4: *iq* normalization in *R*; 5: directLFQ normalization in Python. For strategies 1, 3, 4 and 5, we removed data of ME samples prior to normalization. (B) Box plot analysis of *E.coli* protein intensities ($log_2$-transformed) after MBR and 100x DIA-ME analysis for different data normalizations. Boxes are indicated for all three *E.coli* spiking ratios. Numbers refer to the same strategies as in panel A. Internal peptide normalization of DIA-NN estimates protein intensities greater for DIA-ME analysis (strategies 2 and 3). (C) Mean protein quantity ratios dependent on their abundance after 5x DIA-ME (left panel) und 100x DIA-ME analysis (right panel), illustrated as scatter plot on the left and box plot on the right. Human and *E.coli* proteins are colored in blues and greens, respectively, with the shade of the color representing the quotient between the different spiking ratios. Expected ratios are depicted as dashed lines. The number of quantified human and *E.coli* proteins are indicated.

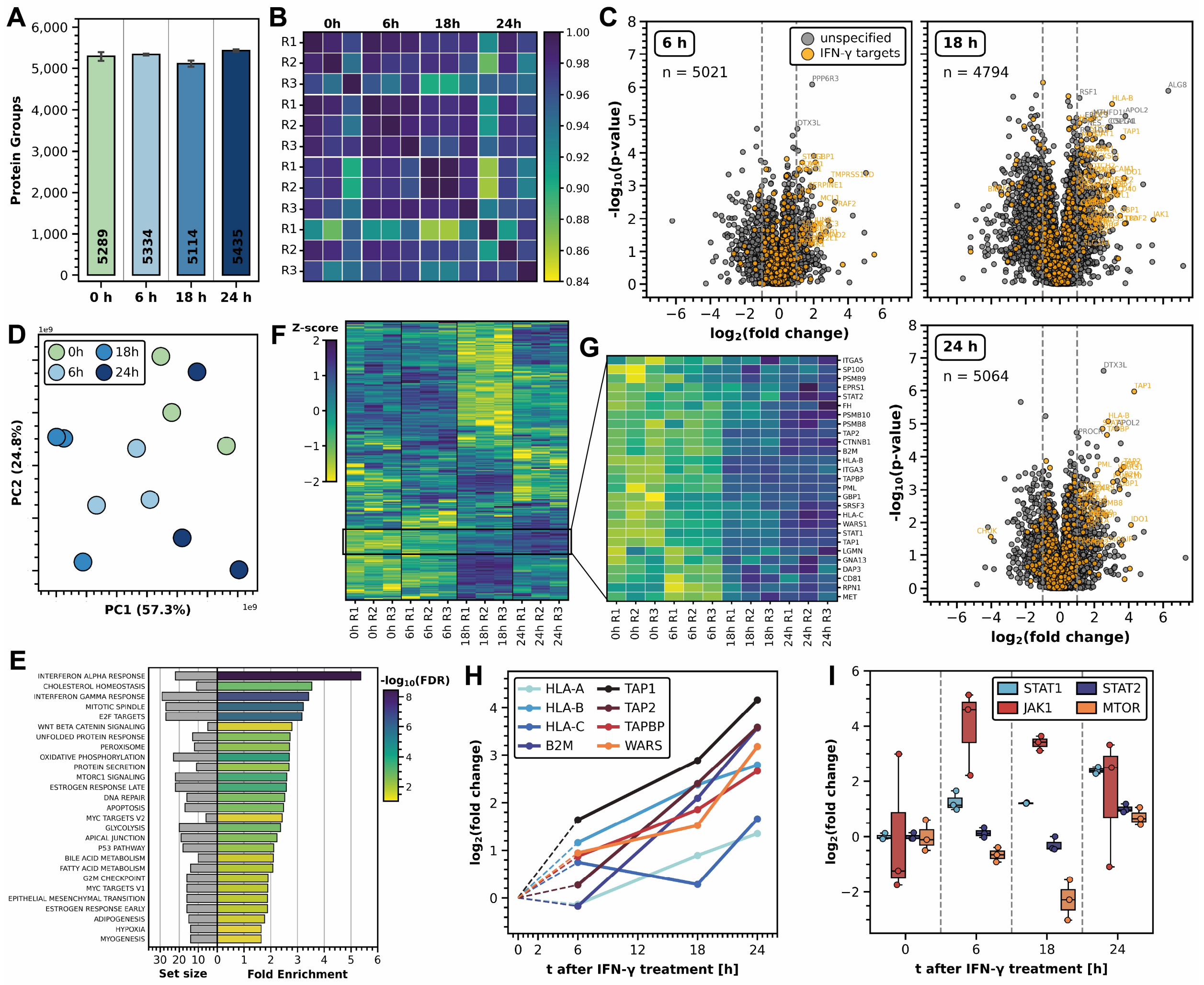


**Figure S7: 200-ng bulk analysis of IFN-γ-treated U-2 OS cells.** (A) Average protein group identifications per time-point (control samples: green, treated samples: blues). (B) Pearson correlation heatmap of time-point replicates. (C) Volcano analysis of Student's *t*-test results between time-point samples 0 hours control sample. n: number of differentially expressed proteins. Proteins known to be involved in the IFN-γ-response were illustrated in yellow. Dashed lines indicate a fold change of 2 and -2. (D) PCA analysis of time-point samples. (E) Gene set enrichment analysis of proteins that showed up-regulation after 24 hours ($log_2$ fold change > 0.58) using MSigDB hallmarks. Bars on the right represent the degree of enrichment, while their colors specify the enrichment's FDR. Bars on the left illustrate the size of the enriched term. (F) Heatmap of known IFN-γ-responsive proteins after hierarchical clustering by Euclidean distance. Colors illustrate the quantitative changes compared to the protein's median across all samples by Z-score. The indicated black box outlines an identified cluster that shows gradual increasing Z-score over time. (G) Enlarged cluster with gradual increasing Z-score from panel F. (H) Line plot of several selected proteins, taken from panel G, showing their gradual up-regulation over the course of treatment. (I) Boxplots of observed fold changes of proteins STAT1, STAT2, JAK1 and mTOR over time.

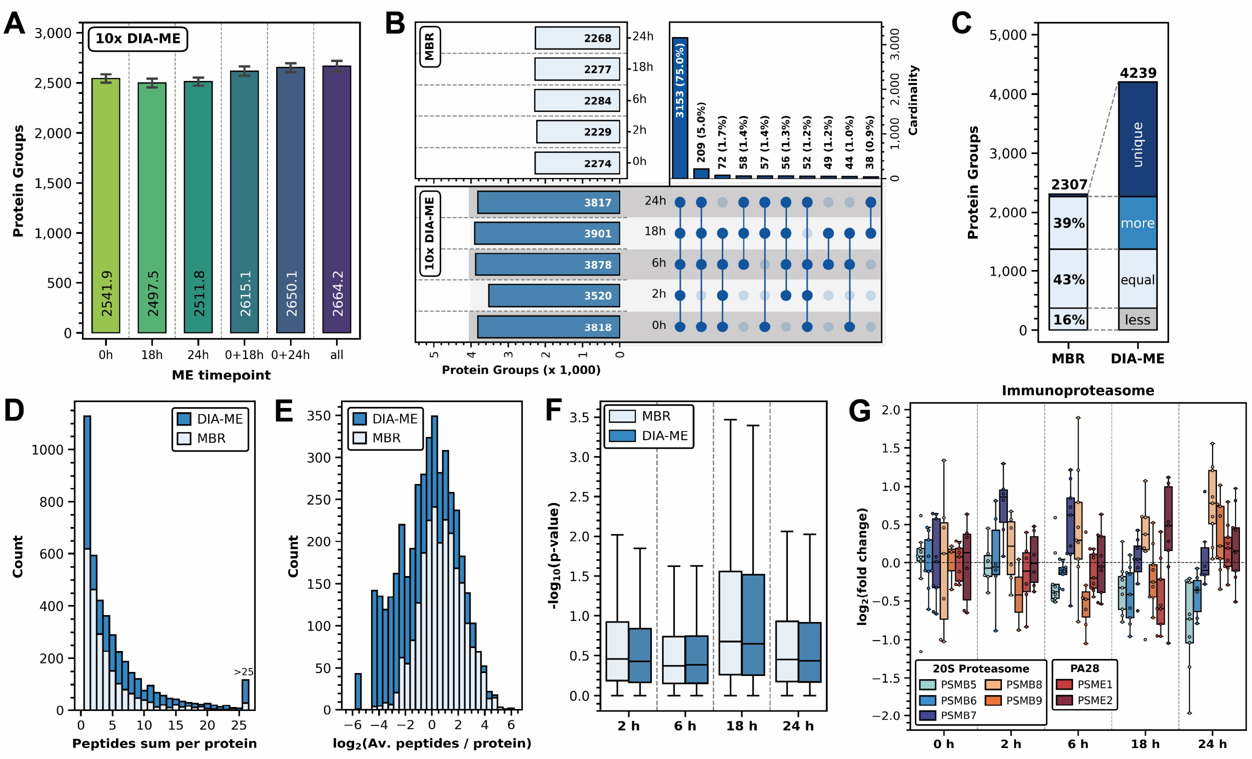


**Figure S8: Different additional analyses of 200-pg samples, complementing figure 5 and 6.** (A) Bar plot of average protein group numbers after 10x DIA-ME analysis involving different ME time-point samples. (B) UpSet plot of 10x DIA-ME analysis showing total protein group numbers per time-point on the left and protein cardinality, i.e. presence in the time-points, on the left. Numbers and proportions of proteins in the respective time-points, shown by the knots, are indicated on top of the bars. Total protein group numbers per time-point after MBR analysis are given as a reference. (C) Bar plot of total protein group identifications in MBR and 10x DIA-ME analysis across time-points. The bars are divided into fractions of proteins (proportions are indicated) that showed more (blue), equal (light blue) or less (light grey) individual identifications in the replicates, i.e. higher, equal or lower data completeness, after DIA-ME analysis. Proteins that were identified exclusively in MBR or DIA-ME analysis (unique) are shown in dark blue. (D) Histogram of total identified peptides per protein for both analyses (white: MBR; blue: DIA-ME). (E) Histogram of average number of peptides per protein and sample ($log_2$-transformed) for MBR (white)

and DIA-ME analysis (blue). (F) Boxplot analysis of p-value distributions per time-point and analysis (white: MBR; blue: DIA-ME). (G) Boxplot analysis of fold changes ($log_2$-transformed) from observable 20S proteasome proteins that define the immunoproteasome and the proteasome activator PA28.

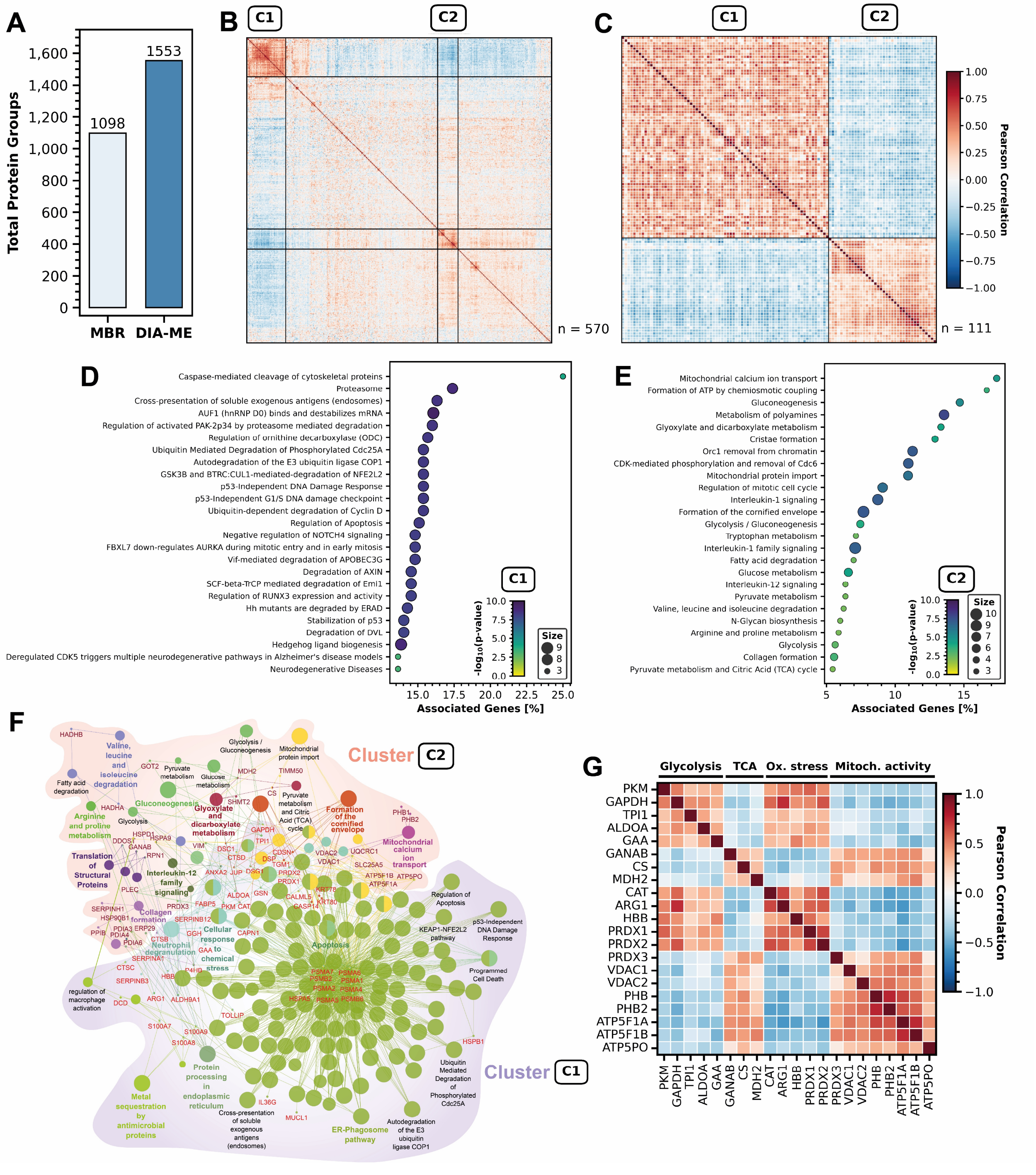


**Figure S9: DIA-ME-assisted identification of metabolic states in single U-2 OS cells.** (A) Total protein group identifications after conventional MBR (light blue) and DIA-ME analysis (blue) using 10-cell ME samples. (B) Co-expression analysis by Pearson correlation of proteins present in ≥ 20 cells. Two clusters showing strong internal correlation but mutual anti-correlation are labelled accordingly. (V) Enlarged version of the two clusters C1 and C2 from panel C. (D) Gene set enrichment analysis of clusters C1 shown as bubble plot. Bubble sizes indicate the size of the enriched term, while their color specifies the enrichment's FDR. (E) Gene set enrichment analysis of cluster C2. (F) Protein-protein interaction and protein-pathway interaction network, showing relations within and between clusters C1 and C2. The nodes of the network represent the terms associated with indicated proteins (red: cluster C1, dark red: cluster C2), and (undirected) edges represent interactions between proteins. Colored nodes highlight significantly enriched pathways and indicate the protein's function, as follows: Blue: signaling proteins, Green: structural proteins, Red: metabolic proteins, etc. (G) Co-expression analysis of a selection of inversely correlated proteins showing the metabolic effect upon oxidative stress and are putative links of both clusters in panel F.

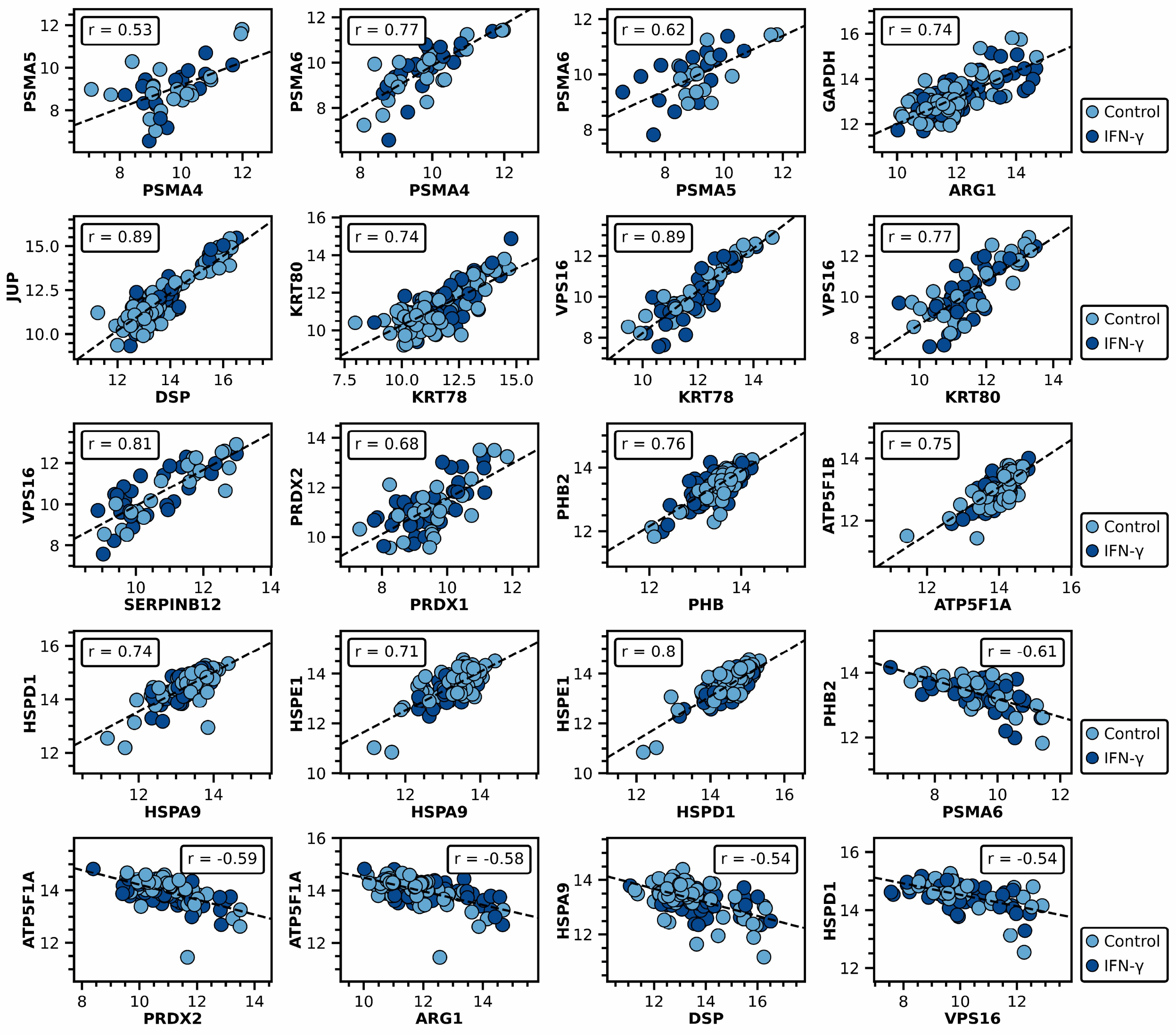


**Figure S10: Examples of pairwise-correlated and anti-correlated proteins.** Scatter plots of protein intensities ($log_2$-transformed) observed in individual cells with protein names indicated at the x- and y-axis. Dark blue: IFN-γ-treated cell; blue: control cell. Linear regressions by Pearson are shown as dashed lines with the respective correlation r indicated.
